# QM/MM Free Energy Calculations of Long-Range Biological Protonation Dynamics by Adaptive and Focused Sampling

**DOI:** 10.1101/2024.04.23.589781

**Authors:** Maximilian C. Pöverlein, Andreas Hulm, Johannes C. B. Dietschreit, Jörg Kussmann, Christian Ochsenfeld, Ville R. I. Kaila

**Author notes:** Corresponding Author To whom correspondence should be addressed: Ville R. I. Kaila.

## Abstract

Water-mediated proton transfer reactions are central for catalytic processes in a wide range of biochemical systems, ranging from biological energy conversion to chemical transformations in the metabolism. Yet, the accurate computational treatment of such complex biochemical reactions is highly challenging and requires the application of multiscale methods, in particular hybrid quantum/classical (QM/MM) approaches combined with free energy simulations. Here we combine the unique exploration power of new advanced sampling methods with density functional theory (DFT)-based QM/MM free energy methods for multiscale simulations of long-range protonation dynamics in biological systems. In this regard, we show that combining multiple walkers/well-tempered metadynamics with an extended-system adaptive biasing force method (MWE), provides a powerful approach for exploration of water-mediated proton transfer reactions in complex biochemical systems. We compare and combine the MWE method also with QM/MM-umbrella sampling and explore the sampling of the free energy landscape with both geometric (linear combination of proton transfer distances) and physical (center of excess charge) reaction coordinates, and show how these affect the convergence of the potential of mean force (PMF) and the activation free energy. We find that the QM/MM-MWE method can efficiently explore both direct and water-mediated proton transfer pathways together with forward and reverse hole transfer mechanisms in the highly complex proton channel of respiratory Complex I, while the QM/MM-US approach shows a systematic convergence of selected long-range proton transfer pathways. In this regard, we show that the PMF along multiple proton transfer pathways is recovered by combining the strengths of both approaches in a QM/MM-MWE/focused US (FUS) scheme, and revealing new mechanistic insight into the proton transfer principles of Complex I. Our findings provide a promising basis for the quantitative multi-scale simulations of long-range proton transfer reactions in biological systems.

## INTRODUCTION

Proton transfer reactions are essential for many biological processes, ranging from catalytic transformations in the metabolism to cellular respiration and photosynthesis, that are responsible for biological energy conversion.^1,2^ Biological proton transfer reactions are often catalyzed by titratable amino acids (His, Lys, Asp, Glu) buried within the protein core that together with water molecules form “proton wires” that facilitate proton transfer via bond-rearrangement in a Grotthuss-type transfer reaction^3^. Respiratory and photosynthetic enzymes employ such water-mediated proton transfer reactions to create a proton motive force across a biological membrane, powering the synthesis of ATP and active transport in cells.^4^ Yet, despite major advances in understanding these complex biological systems, the mechanistic principles of several bioenergetic enzymes remain unclear and highly debated. In this regard, multiscale simulations provide key understanding of how the protonation reactions are controlled by the protein structure and dynamics.^1^ The mechanistic principles of different proton transfer reactions in several bioenergetic systems have indeed been addressed by various multiscale methods in recent years.^5-8^ Yet, long-range biological proton transfer reaction that can extend across large distances still pose major challenges for modern multiscale methods due to a high computational cost, challenges in converging free energy profiles of different reaction pathways, together with the need for the accurate treatment of the electronic structure of the system.

The accurate computational treatment of protonation dynamics requires modeling of both bond-breaking and bond-formation reactions, as well as the conformational dynamics of the system. To lower the computational cost, semi-empirical approaches (*e*.*g*., DFTB, SCC-DFTB, PM7) as well as reactive force field methods (*e*.*g*., EVB, MS-EVB) have been developed. Although these methods can provide valuable insight into various biological reactions,^8-15^ they require pre-parametrization that can be difficult to achieve.^16^ In this regard, density functional theory (DFT)-based methods often provide a good compromise between the computational cost and accuracy for many complex systems,^17, 18^ together with hybrid quantum mechanics/classical mechanics (QM/MM) methods that allow for modeling complex interactions of the biological surroundings at an approximate atomistic force field level.^1, 19^ Since the protonation reactions can involve significant charge re-arrangements that lead to conformational changes in the protein surrounding, they must be sampled by explicit molecular dynamics or enhanced sampling methods on timescales that are often challenging to capture by first principles methods due to the high computational cost.

Recent developments in electronic structure methods, *e*.*g*., pre-screening of integrals and various linear scaling approaches,^20^ can provide a significant reduction of the computational costs, thus allowing modeling of large extended QM systems. Together with new hardware and implementations, such as the accelerations provided by graphics processing units (GPU),^21^ they provide new opportunities for the exploration of longer timescales and rare events.

Here we introduce, implement, and benchmark QM/MM enhanced sampling methods in combination with different reaction coordinates for simulations of long-range proton transfer reactions. More specifically, we study the recent shared-bias well-tempered metady-namics-extended adaptive biasing force (MWE) method,^22^ and both compare and combine this with umbrella sampling (US) in the context of multiscale QM/MM simulations. We study the performance of geometric (linear combination of bond-breaking and bond-formation process) and physically motivated (modified center-of-excess charge, mCEC^23^) representations of the transferred proton as reaction coordinates/collective variables (CVs). The former CV is defined *a priori* by manual selection of the involved bond distances, while the global nature of the latter aims for a unified description of all possible proton transfers, which may open up the exploration of new mechanisms *on-the-fly*.

We show how the unconfined diffusion along CVs in the MWE approach can accelerate the conformational sampling and exploration of various reaction mechanisms relative to the US approach, which by construction, confines the simulations to a single reaction channel. However, we also discuss challenges in obtaining converged free energy profiles using MWE due to the sampling of a larger conformational space, which may require combination of multiple simulations and manual tuning of the sampling parameters.

To address the convergence of such challenging PMFs, we combine the QM/MM-MWE and QM/MM-US simulation methods to systematically recover the multidimensional PMF along different reaction mechanisms. In the following, after a short theoretical review of the US and MWE methods, we discuss the performance of both sampling strategies on a model system. Finally, we apply the presented framework to the protonation dynamics of respiratory Complex I, a highly challenging biological system, where the different sampling strategies are explored.

## THEORY, METHODS, AND MODELS

The potential of mean force (PMF) along a reaction coordinate or collective variable (CV) is defined as,

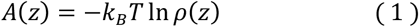

with the Boltzmann constant *k*_*B*_, the temperature *T*, and the probability density function *ρ*(*z*) of finding the system in a certain state z along the CV *ξ*(***x***), defined by

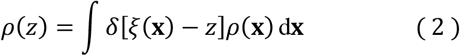

where *δ* denotes the Dirac delta function. The efficient estimation of *ρ*(*z)* often requires application of importance sampling strategies.^24-27^ The US simulations aim to achieve a uniform exploration of the PMF by performing several equilibrium simulations with pre-defined restraints *B*(*ξ*(***x***)), often modeled as harmonic biasing potentials. Similarly, in the MWE method, the CVs are coupled to harmonic potentials *B*(*ξ*(***x***),*λ*), whilst the diffusion of the simulation windows along the CV is achieved by the coupling to a fictitious particle *λ*.^28^ The full potential energy of the extended system (*x,λ*) can be defined as,

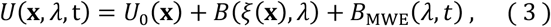

where *U*_0_(**x**) is the potential energy of the system. To ensure uniform sampling of the CV along *λ*, a time-dependent bias potential *B*_MWE_(*λ,t*) can be added. In this regard, the MWE uses two complementary strategies, simultaneously filling the free energy wells (well-tempered metadynamics) and removing the barriers (extended adaptive biasing force). Post-processing allows recovering the restrained simulation windows *B*_*i*_(*ξ*(***x***)) analogous to US windows by separation of the extended system into states with constant *λ* = *λ*_*i*_.^29^

As the biasing potentials are known, the unbiased probabilities are obtained *via*,

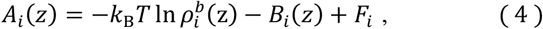

where the integration constants, *F*_i_, for each window *i* describe the vertical position of the individual unbiased PMF, while 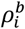 are the probability distributions obtained from the biased runs. In practice, the unbiased PMF is obtained by the weighted-histogram analysis method (WHAM) or by the multistate Bennett’s acceptance ratio (MBAR), a histogram free/zero bin width version of WHAM.^27, 30^ The unbiased probabilities in MBAR are obtained by,

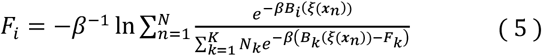

with *K* simulation windows comprising *N*_*k*_ samples and data points from the pool of all simulations ***x***_***n***_. The unbiased weights of the individual frames can thus be recovered as,

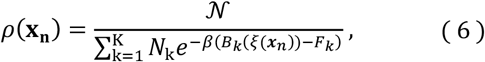

where the normalization constant 𝒩 is introduced to ensure that Σ_*n*_ *p*(***x***_***n***_) = 1. Recent developments, such as the transition-based re-weighting (TRAM,^31^ dTRAM^32^) and the dynamic histogram analysis extended to detailed balance (DHAM,^33^ DHAMed^34^), which account for the transitions between the restrained simulation windows, can also be used to obtain the free energy estimate *F*_*i*_ as a function of the CV.

Accurate activation free energies, which define the reaction rates, can be determined from the PMF by using a definition yielding consistent energies for CVs with parallel gradients,^35^

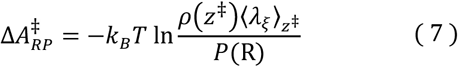

where *ρ*(*z*^‡^) is the probability density at the dividing surface, *P*(R) is the probability that the system resides in the reactant state (obtained by integration over the PMF), and ⟨*λ*_*ξ*_⟩_*z*_ is the conditional average of the de Broglie wavelength,^35^ related to the mass of the pseudo particle associated with the CV, *m*_*ξ*_ (see Extended Methods). A comparison of this expression with the often-employed harmonic approximation is given in the SI. Eqn. (7) removes possible distortions of the Cartesian space by non-linear CVs and accounts for the mass of the atoms involved in the transition, whilst apparent free energy barriers obtained from difference between maxima and minima on the PMF 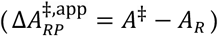 does not include such corrections. The former treatment correctly reproduces, *e*.*g*., isotope effects.^35^

### Computational methods

All QM regions were described at the B3LYP-D3/def2-SVP level of theory,^36-39^ which has shown a good balance between computational cost and accuracy for many biological systems (see Figure S15, and Ref. ^40^ for detailed benchmarking).^6, 41-46^ The currently employed density functional level is likely to underestimate barriers by a few kcal mol^-1^ relative to *ab initio* theory (Figure S15, see also Ref. ^47, 48^). The surroundings of Model 1 were described with COSMO at ε=4,^49^ while those of Model 2 were described explicitly (see below for details on Model 1/2). MD simulations were performed at *T*=310 K with a timestep of 0.5 fs. PMFs were calculated using MBAR,^30, 50^ as described in Ref ^29^. For the QM/MM sampling, activation free energies 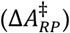 were obtained based on the *z*-averaged inverse mass of the reaction coordinate.^35^ All simulations were implemented in Python and performed with the GPU-accelerated QM code Fermi-ONs++^20, 21, 51-53^ coupled to OpenMM^54^ *via* a QM/MM interface. Significant speedup of QM(/MM)-MD calculations on GPUs were achieved by accelerating evaluations of exact-exchange with the sn-LinK method^52, 55^ and by using the RI-J approximation of the Coulomb energy.^56^ To accelerate the sampling, we further applied the multiple walkers (MW)/shared bias approach, where parallel simulations synchronize the time-dependent bias potential *U*_MWE_(λ,*t*) in regular time intervals. Initial system setup and minimization of the reaction pathways were performed using TURBOMOLE v. 7.4.^57, 58^ Visual Molecular Dynamics (VMD)^59^ and PyMOL^60^ were used for visualization.

### Model 1: Model system for water-mediated proton transfer

A system containing two histidine residues (modeled as methyl imidazole), hydrogen-bonded by two water molecules, was employed as a model system to study the free energy profile of proton transfer reactions. To this end, we studied the performance of two different reaction coordinates (Figure 1A,D): I) a geometric reaction coordinate, defined as a linear combination (LC) of bond breaking and bond formation distances (Figure 1A),

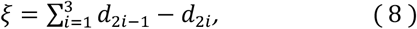

and II) a modified center of excess charge (mCEC, see also Figure 1D),

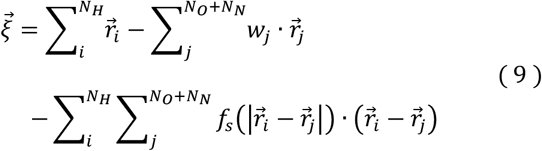

with the projection onto the donor-acceptor vector,

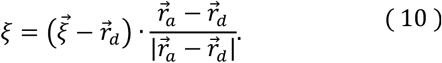

The delta nitrogen (Nδ) of the first histidine was defined as the proton donor (*d*), while the Nδ of the second histidine residue was defined as the proton acceptor (*a*) (Figure 1D). *N*_H_ denotes the number of involved exchangeable protons, *N*_O_ and *N*_N_ the number of oxygen and nitrogen proton acceptors. The weights, *w*_j_, were set to *w*_N_=0 and *w*_O_=2, which describe the minimum number of protons bound to atom *j* in the reactant or product state. The modification term *f*_*s*_(*d*_ab_) introduces a logistic function to provide a smooth switching between the cross-terms and is defined as,

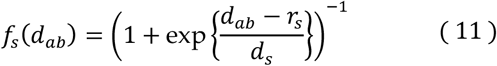

where *d*_ab_ is the distance between a heavy atom *a* and proton *b*, both participating in the reaction coordinate. The parameters *r*_s_ and *d*_s_, define the location and width of the switching regime and were set to 1.3 Å and 0.05 Å, respectively, unless stated otherwise (Figure S6).

**Figure 1.**
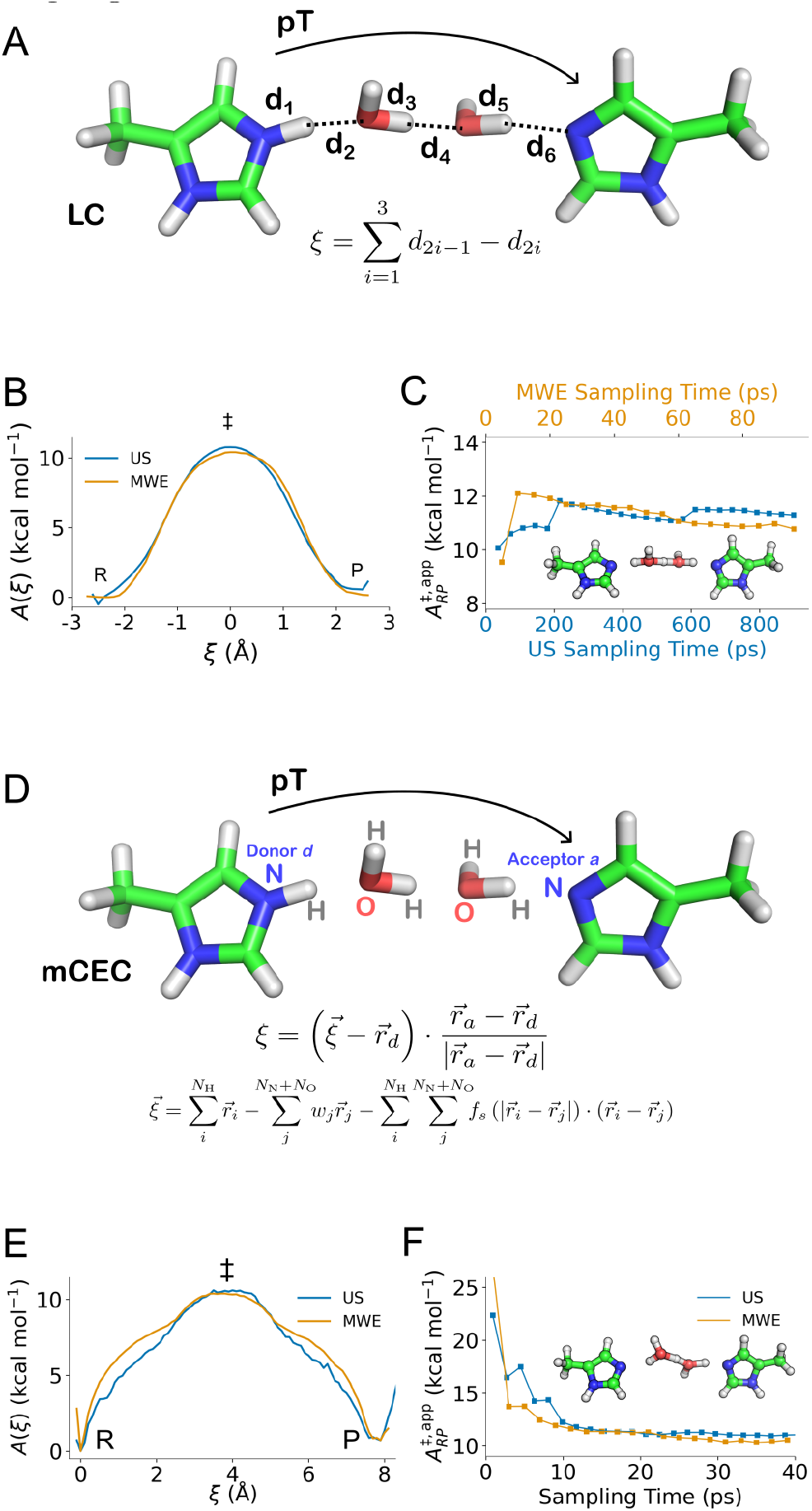
Sampling of water-mediated proton transfer reaction in a His/water model system with QM/MM umbrella sampling (US) and the multiple walker/well-tempered metadynamics extended-system adaptive biasing force (MWE) method. The PMF was explored using a linear combination (LC, panel A) of geometric distance and the modified center of excess charge (mCEC, panel D) as reaction coordinates. (A, D) Definition of the reaction coordinates. (B, E) The PMFs obtained using the LC and mCEC definitions with US and MWE. R, reactant, P, product, and ‡ transition state region. (C, F) Convergence of the max-min difference with increasing sampling time. Representative conformations sampled in the transition state region are shown as an inset.

Initial coordinates were obtained by geometry optimization at the B3LYP-D3/def2-SVP level,^36-39^ while keeping the distance between the C_β_ atoms of the histidines fixed to 15 Å. The US simulations were performed using 18 windows sampled for 50 ps with a harmonic bias [B_i_=1/2 *k* (*z* - *z*_i_)^2^] centered at LC coordinates [-2.07 Å, +2.18 Å] every 0.25 Å with a force constant of *k*=100 kcal mol^-1^ Å^-2^. The MWE simulations were carried out using two walkers, each sampled for 48 ps. The extended variable was coupled to the system with a coupling width σ=0.1 Å and adaptive forces were accumulated on a grid with a bin width of Δ=0.05 Å. Well-tempered metadynamics was applied to the extended system with a Gaussian hill width of 0.1 Å and a hill height of 0.12 kcal mol^-1^ (0.5 kJ mol^-1^) added every 10^th^ fs. The bias factor *γ*=*ΔT*/*(T*+*ΔT)*, which ensured a smooth hill decay, was set to 0.866.

To explore the mCEC coordinate, US simulations were performed with 33 equally spaced windows, each simulated for 20 ps with a harmonic bias placed on the mCEC reaction coordinate, between [0 Å, 8.75 Å] every 0.25 Å, and a force constant of *k*=100 kcal mol^-1^ Å^-2^. MWE simulations along the mCEC were performed with two walkers, each sampled for 50 ps. To this end, the extended variable was coupled to the system with σ=0.1 Å and adaptive forces were accumulated on a grid with Δ=0.1 Å. In the MWE, well-tempered metadynamics was applied to the extended system with a Gaussian hill width of 0.2 Å, a hill height of 0.239 kcal mol^-1^ (1 kJ mol^-1^) and a frequency of hill creation 10 fs, using a bias factor of γ=0.866. Additional convergence tests of the MWE sampling were performed using 10 walkers each sampled for 50 ps at the semi-empirical GFN2-xTB level.^61, 62^

### Model 2: Water-mediated proton transfer reactions in Complex I

The proton channel in Complex I was modeled based on the cryoEM structure of the mouse enzyme (PDB ID: 6ZTQ^63^) that was embedded in a POPC/POPE/cardiolipin (2:2:1) membrane and solvated with water molecules and NaCl (150 mM). The system was relaxed over 1 μs with classical MD simulations, followed by construction of the QM/MM model system. The QM region, comprising 114 atoms, included Lys237, His319, His293, Glu378, Ser289, Ser290, Ser323, Asn366, Asn374 of subunit ND4, as well as twelve water molecules (Figure 3A, inset ii). The QM region was coupled to the MM system via the link atom approach, introduced between the C_α_ and C_β_ bonds. The surroundings were described with the CHARMM36 force field,^64^ and the QM-MM interaction via an additive electrostatic embedding scheme. The QM/MM system was geometry optimized at B3LYP-D3/def2-SVP level prior to QM/MM free energy sampling.

**Figure 2.**
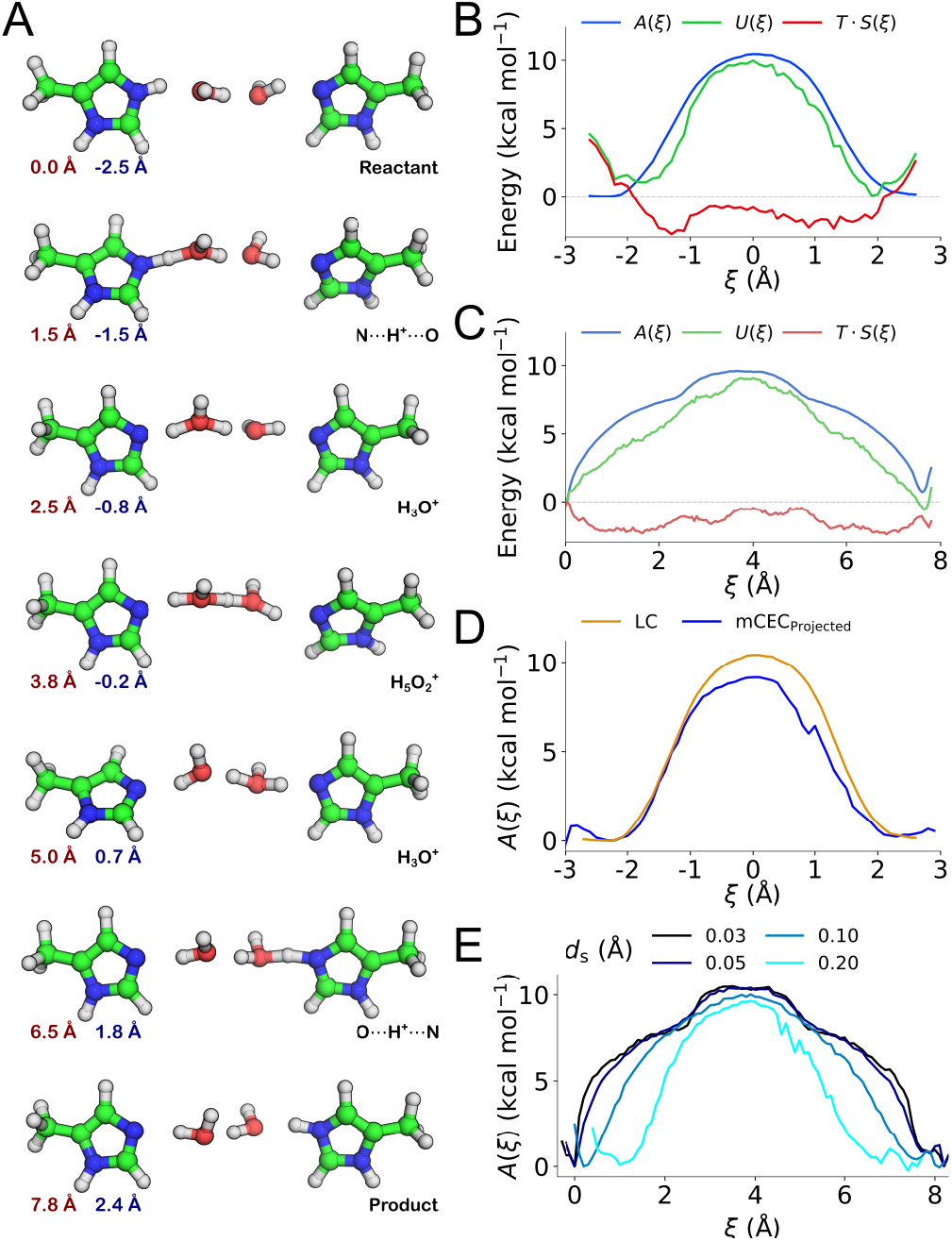
(A) Representative intermediate geometries extracted from the mCEC/MWE sampling with corresponding mCEC (red) and LC (blue) values. (B,C) Free energy *A*, internal energy *U*, and entropy *T·S* profile for the His-water array, sampled with MWE as a function of the LC coordinate and mCEC coordinate, respectively. (D) PMF sampled with mCEC projected onto the modified LC in comparison with LC-sampled data, also shown in Figure 1B. (E) PMF profiles obtained from mCEC/MWE sampled data mapped onto the mCEC definitions with different *d*_s_ values.

**Figure 3.**
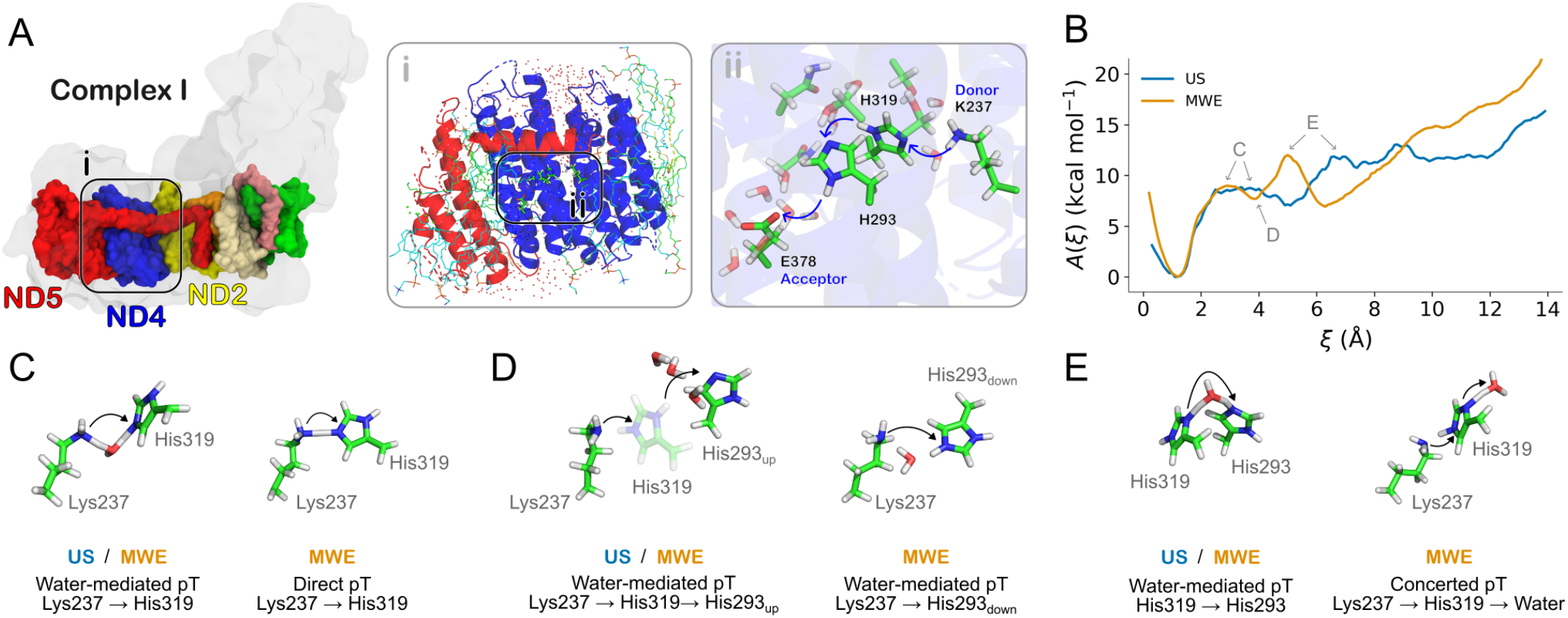
Free energy sampling of proton transfer reactions in the membrane domain of respiratory Complex I (subunit ND4). (A) The mouse Complex I with highlighted core subunits of the membrane domain. *Inset* i: Overview of the QM/MM system consisting of ND4 and its environment. *Inset* ii: The QM region with central residues participating in the proton transfer reaction. Blue arrows indicate the proton pathways sampled in the QM/MM free energy calculations. (B) PMFs of the proton transfer reaction obtained with the US and MWE methods. (C,D,E) Representative conformations for the CV values indicated in panel B, with US (in blue) and MWE (in orange). Subscripts refer to species described in the main text. Due to the explorative sampling by MWE, two representative conformations are shown for the selected states. Additional conformations are shown in Figure S8.

The water-mediated proton transfer from Lys237 to Glu378 was modeled using the mCEC reaction coordinate with *r*_*s*_ and *d*_*s*_ set to 1.3 Å and 0.03 Å, respectively, accounting for 27 protons on 17 possible donor/acceptor atoms and including also the 12 nearby water molecules. The US windows were placed between [0.5 Å, 13.75 Å] every 0.25 Å, with a force constant of 100 kcal mol^-1^ Å^-2^, with additional windows placed in regions of low overlap, resulting in a total simulation time of 300 ps.

The QM/MM-MWE simulations were carried out with a bin width and coupling width of 0.1 Å each with twelve walkers, each sampled for 25 ps. For the well-tempered metadynamics new Gaussians were added every 10^th^ fs with a bias factor *γ*=0.866, a hill height of 0.096 kcal mol^-1^ (0.4 kJ mol^-1^), and hill width of 0.2 Å. Additional focused US simulations were performed starting from manually selected snapshots of the MWE simulations in chemically interesting and undersampled regions (cf. Figure 4A). To this end, windows were confined to the initial mCEC value of the selected snapshots with a harmonic bias and a force constant of *k*=100 kcal mol^-1^.

**Figure 4.**
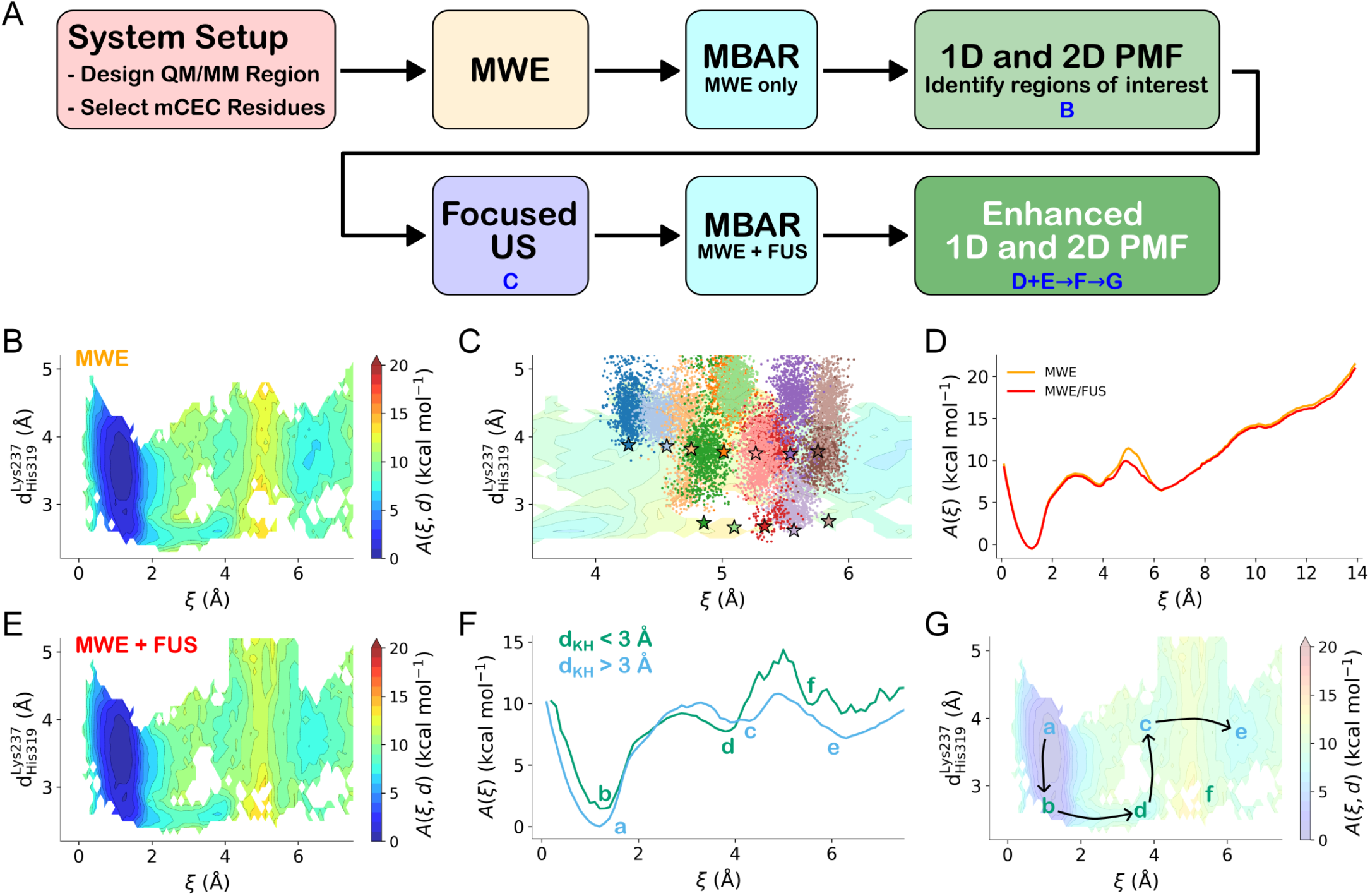
Hybrid MWE / focused US free energy sampling scheme of proton transfer reactions in the membrane domain of Complex I (subunit ND4). (A) Suggested workflow of the MWE/FUS approach. B-G) Analysis steps as indicated in the flow chart, based on simulation results of Complex I (Model 2). (B) Two-dimensional PMF from MWE trajectories. The two dimensions are the mCEC CV and the distance between Lys237(Nζ) – His319(Nδ) distance (here *d*_KH_). (C) 2D-PMF overlaid with initial reaction coordinates of focused US windows and their trajectories. (D) 1D-PMF along the mCEC coordinate before and after addition of the focused sampling. (E) 2D-PMF of combined data from MWE and focused US. (F) PMFs determined for *d*_KH_ < 3 Å (green) and *d*_KH_ > 3 Å (cyan). Minima are indicated by *a-f*. (G) 2D-PMF with minima *a*-*f* indicated. The arrows denote possible reaction mechanism along the *a* to *e* pathway.

Simulations of the proton transfer reaction were also performed with a smaller QM region, comprising 65 atoms (Lys237, His319, His293, and E378, together with six water molecules) to speed up the sampling. The QM region was restrained with positional restraints using a harmonic force constant of 23.9 kcal mol^-1^ Å^-2^ (100 kJ mol^-1^ Å^-2^) placed on the heavy atoms to limit the sampling along water-mediated pathways in both the US and single walker/well-tempered metadynamics extended-system adaptive biasing force (SWE) simulations. In the SWEs, a coupling width of 0.1 Å was employed. A conformational search of the system was additionally carried out using the conformer-rotamer ensemble searching tool (CREST, see Extended Methods for further details).^65^

Convergence of the PMF profiles was assessed by monitoring barrier heights 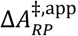 as well as free energy profile differences 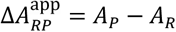 function of the sampling time.

## RESULTS

### Potential of mean force profiles for water-mediated proton transfer along a histidine-water array

To test and compare the performance of US and MWE sampling, we first studied a model system (Model 1) of a water-mediated proton transfer reaction using a quasi-one-dimensional water wire connecting two histidine residues (Figure 1). Similar and related model systems have previously been used to explore mechanisms of proton transfer reactions.^11, 66-68^ The sampling of the proton transfer reaction was performed using i) the LC (Eqn. (8), Figure 1A) and ii) the mCEC (Eqn. (9), (10), Figure 1D) reaction coordinates. Combining these reaction coordinates with both sampling methods (US and MWE) resulted in four conditions tested for Model 1 (Simulations 1-4, see Table S1). The reaction coordinate space was well-sampled and yielded converged PMF profiles for all models (see Figure S1,S2). Convergence was also probed at the semi-empirical GFN2-xTB level of theory, which support that the PMF and the apparent free energy barrier converge around 300 ps, in good agreement with the DFT-based sampling (see Figure S2G-I).

Comparison of the sampling methods shows that the MWE method outperforms the US method by reaching convergence in a shorter simulation time. While the US requires around 200 ps of sampling to reach full convergence along the LC reaction coordinate (Figure 1C), we obtain a statistically converged PMF in around 40 ps with MWE. Moreover, for the mCEC coordinate, the convergence with MWE is twice as fast as with US. However, we also note that the apparent free energy barrier 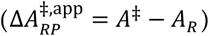 (Figure 1C,F) changes by <5% (0.5 kcal mol^-1^) after extending the sampling beyond 20 ps per window with US. Both sampling methods predict a transition state that comprises a Zundel ion (H_5_O_2_^+^) hydrogen-bonded to both histidine residues, with a similar conformational space explored by both approaches (Figure S3-S5). Both methods also predict activation free energies 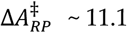 kcal mol^-1^ along the LC coordinate, and 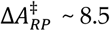 kcal mol^-1^ along the mCEC coordinate. This shift could arise from sampling differences in the transitions along the LC and mCEC, although we also observe differences in the predicted entropy profile (Figure 2B,C).

We note that the shape of the PMF is somewhat different along the CVs, which could affect, *e*.*g*., the prediction of mean-free passage times.^69, 70^ Along the LC coordinate, we obtain a PMF profile resembling a Gaussian hill, whereas for the mCEC, the PMF profile has a double well shape with a broad maximum in the transition state region and steep rise near the reactant and product minima. The saddle points along the MWE profile correspond to configurations where the proton is shared between the histidine and the water molecule (Figure 1E). The MWE minima are also flatter relative to those obtained using US along the LC coordinate, an effect that could arise from the enhanced rotation of the imidazole-ring in the MWE sampling.

Projection of the mCEC sampled conformations onto the LC reaction coordinate suggests that the latter does not uniquely cluster the conformations into reactant and product states. In this regard, we note that sampling along the mCEC coordinate leads to an exchange of the “off-pathway” protons (*i*.*e*., protons not directly involved in the Grotthuss wire) with the “on-pathway” protons (Figure S3B, S4A). Such an exchange improves the estimation of entropic effects that are not easily captured when the sampling is performed along the LC coordinate, and may in turn lead to an overestimation of the activation free energy (see above).

By accounting for the protons considered for each sampled conformation, we find that a projection of the mCEC conformations onto the LC reaction coordinate can be achieved in *post hoc* analysis. By applying this projection procedure, we recover both the separation between reactant and product regions that re-maps the LC coordinate onto the physically relevant range between [-3 Å, +3 Å] (Figure S4B). Moreover, the resulting PMF shows a clear separation between reactant and product minima and the characteristic double-well shape (Figure 2D). However, we note that the profiles still differ in their apparent barrier height, an effect that could arise from sampling differences. To further test the origin of these differences, we computed entropy and internal energy profiles, according to Ref. ^71^, by extending the sampling to 300 ps (Simulation 5, see Table S1). For the mCEC coordinate, we observe three peaks in the entropy profile at *ξ*=2.5, 3.8, and 5.0 Å (Figure 2C) that arise from the increased combinatorial sampling of atoms in the protonated water species. The four minima on the entropy curve correspond to the restricted configuration space when bonds are broken or formed, as only few orientations are energetically feasible during the proton transfer reaction. Analysis of the O-O distances and the O / H positions reveals that the entropy maxima correspond to hydronium (H_3_O^+^) and Zundel (H_5_O_2_^+^) species (Figure 2A, Figure S3A). The Zundel ion, together with the lateral motion of both O atoms (Figure S3C,D), suggest that the process follows a semi-concerted, Grotthuss-like mechanism, with a subtle diffusive component, possibly arising from the constrained His-His distance. Interestingly, the internal energy profile peaks around ξ=4.0 Å, with the monotonous decreases towards the reactant and product regions, suggesting that the Zundel ion is energetically unstable (Figure 2C). In contrast, for the LC coordinate the entropy profile shows only one maximum (Figure 2B), as by construction, the PMF along the LC is a superposition of all bond formation and bond breaking reactions. Therefore, configurations that comprise Zundel or hydronium ions are mapped to the same LC value around 0 Å. The LC may thus favor a concerted mechanism, while the mCEC would in principle also allow for sequential proton transfers, an aspect that could also contribute to the better estimation of entropic effects along the mCEC coordinate.

We also probed the character of the modification term (Eqn. (9), third term) in the mCEC by mapping the mCEC/MWE data onto the mCEC definitions by testing the effect of different values of the switching width *d*_*s*_ (see Eqn. (11)). With an increase in this parameter, we observe a change in the shape of the PMF profile, an increase in the mass of the quasi-particle associated with the CV, and a noticeable decrease of the apparent barrier 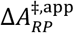 from 10.5 kcal mol^-1^ to 9.6 kcal mol^-1^ (Figure 2E, SI Table 1). By increasing *d*_s_, the switching function becomes smoother, which further affects *ξ, m*_*ξ*_, as well as the shape of *A(ξ)* (Figure S6). In contrast, we find that the activation free energy 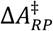 is independent of the switching width parameter and remains at 8.5 kcal mol^−1^.

### Protonation dynamics in respiratory Complex I

We next probed the performance of both QM/MM-MWE and QM/MM-US methods on the lateral proton transfer reaction in the membrane domain of respiratory Complex I (Model 2). This system is central for understanding biochemical processes underlying cellular respiration. Its mechanistic principles remain elusive and highly debated despite significant efforts over the last decade^1, 72-74^. To this end, we explored the proton transfer reaction in a conformational state where the proton transfer reaction is endergonic (“closed ion-pair form”, see Refs. ^5, 43^ for further details). Moreover, we chose to investigate the proton transfer reaction along the mCEC coordinate due to the enhanced sampling properties obtained for Model 1. In this regard, we modeled the proton transfer along the 14 Å water-mediated hydrogen-bonded wire connecting the proton donor Lys237 *via* His319 and His293 with the proton acceptor Glu378 (Figure 3A).

Both the QM/MM-US and QM/MM-MWE simulations (Simulations 6 and 7, see Table S1) capture the uphill PMF for the proton transfer from Lys237 (*ξ*<1 Å) to Glu378 (*ξ*>10 Å), favoring the protonated form of Lys237 by around 10 to 12 kcal mol^-1^ (Figure 3B, Figure S7), but with some differences in the shape of the PMF and relative barriers predicted by the two methods. We find that both methods capture the free energy minima for all intermediate states featuring the protonation of Lys237 (*ξ*=1 Å), His319 (*ξ*=5 Å), His293 (*ξ*=8 Å), and Glu378 (*ξ*=12 Å). Moreover, both PMF profiles suggest that the initial water-mediated proton transfer from Lys237 to His319 has a 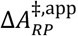 of *ca*. 9 kcal mol^-1^, while the barrier between His319 and His293 is *ca*. 5 kcal mol^-1^ and shifted towards lower CV value for the MWE PMF (*ξ*=5 Å *vs. ξ*=7 Å for US). The last proton transfer step from His293 to Glu378 is isoenergetic in the US PMF and has an apparent barrier of 2 kcal mol^-1^, while the MWE PMF suggests that the step is endergonic with an apparent barrier of 8 kcal mol^-1^.

To understand the differences in the PMF profiles, we compared the conformations, as well as protonation and hydration states. In general, we observe that MWE samples a wider range of states (see below), while the protonation probabilities along the CVs are overall similar for both methods (Figure S8), despite somewhat different protonation profiles around the two intervening histidine residues (Figure S8A,B). In this regard, the saddle point regions comprise Zundel-(H_5_O_2_^+^) and Eigen-(H_3_O^+^)-like species that can be extracted from the US and MWE after clustering the simulation trajectories (Figure 3C-E, Figure S9). Essential dynamics analysis also shows differences in the conformational sampling (Figure S10).

Interestingly, for the initial proton transfer step between Lys237 and His319, the MWE method samples water-mediated conformations (Figure 3C, *left*), as well as conformations where Lys237 directly donates a proton to His319 (Figure 3C, *right*). As a consequence, the distance between Lys237 and His319 is reduced, which could explain the shift of the free energy minimum linked to protonation of His319 towards lower values along the reaction coordinate (Figure 3B). Both methods sample a large conformational change featuring a rotation of the His293 ring towards Lys237 (“downward conformation”, see Figure 3D for definition) that involves re-arrangements of sidechains and water molecules. However, with US, this conformation is only sampled for *ξ*=12 Å, whereas MWE additionally explores such conformation in the *ξ*=3-6 Å regime (Figure 3D, Figure S9), opening an alternative proton transfer pathway, in which His293 could serve as the first intermediate proton acceptor, and thus bypassing His319. These findings suggest that the proton transfer reaction involves conformational changes in His293, which could bridge gaps in the proton wire in Complex I isoforms where His319 is not present.^5, 43^

The conformational changes in His293 are further supported by a conformer search conducted using conformer-rotamer ensemble sampling (CREST) (see Extended Methods, Figure S11).^65^ Other alternative conformations sampled by MWE involve a hole transfer, comprising an initial exchange of a proton between the two His residues, and leading to the His^-^ / His^+^ configuration, which is then followed by proton transfer from Lys237 to His^-^ (on His319) and from HisH^+^ (on His293) to Glu378 (Figure S9). Although energetically feasible in the sampling, it remains unclear if these states are physically realistic or result, *e*.*g*., from DFT charge transfer artifacts.^17^

In general, we note that in addition to the forward proton transfer and reverse hole transfer pathways, the MWE method samples both direct proton transfer between protein residues and water-mediated proton pathways (see above and Figures S9, S12, S13). Microsecond classical MD indeed support that the proton pathways in Complex I undergo changes in the overall hydration levels that further affect the free energy of proton transfer, and in turn provide possible gating principles (*cf*. Figure S8 of Ref. ^5^). In this regard, the conformational space sampled at the MWE level provides a clear benefit in exploring such alternative reaction mechanisms. However, we note that the direct proton pathways could also arise from the changing biasing force applied to the fictitious particle that may lead to a partial displacement of the intervening water molecule.

To facilitate a better comparison between QM/MM-US and QM/MM-MWE, both sampling methods were constrained to the same reaction mechanism by positionally restraining the heavy atoms participating in the reaction (Simulations 9 and 10, Table S1). This constraint is by construction unphysical, but necessary for the direct comparison. As expected, we obtain highly similar PMF profiles for both methods with the constrained sampling (Figure S14). However, the constraints as well as the smaller QM region used to enhance the sampling, results in a more endergonic reaction as compared to the unrestrained sampling.

### Combining the advantages of MWE and US – the hybrid MWE/focused US approach

The MWE approach shows a powerful exploration of the reaction phase space, while US has the advantage to systematically show convergence of a given reaction path. We therefore suggest to combine MWE with a focused US to enhance the sampling of phase space regions of special interest or poorly sampled regions. This hybrid MWE/focused US scheme (MWE/FUS, Figure 4A) could improve the accuracy of the obtained PMF for complex systems. To this end, the workflow could involve the following steps, applicable for a biological proton transfer reaction similar to those studied here: I) System setup, where the QM/MM model is prepared and the residues participating in the mCEC reaction coordinate are selected; II) An extensive QM/MM-MWE sampling with multiple walkers; III) Clustering of sampled conformations by MBAR to derive Boltzmann weights of the microstates; and IV) Compute an initial 1D-PMF based on the Boltzmann weights and visual inspection to identify further degrees of freedom that are relevant for the process, *e*.*g*., characterization of distances between residues or dihedral angles that change during the sampling. These additional degrees of freedom can be used to create 2D-PMFs, in which poorly sampled regions of the phase space are identified. V) Based on the MWE conformations of poorly sampled regions, perform focused US simulations; VI) Derive a refined PMF using MBAR that combines the data from MWE and focused US; VII) Compute new weights that are used to derive refined 1D- and 2D-PMFs. The additional sampling from US improves the reliability of the PMFs, as suggested by sampling the different proton transfer pathways in Complex I (Figure 4E,F). To illustrate the approach, we next applied the MWE/FUS scheme to study the proton transfer step from His319 to His293, where we observed distinct conformations of Lys237 with His319. This region could be of particular interest as the water-mediated Lys237-His319 connectivity seems to lower the barrier for the His319-His293 reaction relative to the direct proton transfer between Lys237 and His319 (Figure 4B,E). To this end, we initialized five/seven US windows (along low/high *d*_KH_ values, lysine-histidine distance) for the two respective connectivity states (Figure 4C, Simulation 8, see Table S1). The Lys237-His319 distance was not constrained to provide an unbiased sampling of the region (Figure 4D).

We find that the focused US indeed improves the 2D-PMF along this region (Figure 4E) and allows the derivation of conformation-dependent 1D-PMFs for the initial reaction steps (Figure 4F). The resulting 2D-PMF reveals that long Lys237-His319 distances are slightly preferred when Lys237 is protonated. Moreover, the apparent barrier is lowered by 1.5 kcal mol^-1^ when the two residues come in close contact, while deprotonation of Lys237 results in longer His-Lys conformations. We therefore propose that the initial proton transfer between Lys237 and His319 could occur by either water-mediated or by direct contact, followed by breaking of the contact and proton transfer from His319 to His293 (Figure 4G). The histidine residues show additional benefits as proton conductors as they can exchange protons (*e*.*g*., with Nδ as acceptor and Nε as donor), but also show a high conformational flexibility.

## DISCUSSION

Our findings suggest that combining accelerated sampling provided by the MWE method with the generalization of the proton transfer reaction through the mCEC reaction coordinate yields a powerful tool for QM/MM free energy calculations of complex (bio)chemical reactions. For our model system of water-mediated proton transfer, we found that the PMF converges faster at the MWE level as compared to US, whereas in the highly intricate proton wires of Complex I, the MWE showed a slower convergence due to the significantly larger conformational space explored. In contrast, QM/MM-US showed a systematic convergence for a given reaction pathway and allowed for an easier parallelization. We also found that by combination of both approaches, the QM/MM-MWE/FUS scheme improves both sampling and accuracy relative to MWE and US (see below).

In this regard, we found that the description of the long-range proton transfer reactions with the mCEC reaction coordinate provides an improved exploration of different regions of conformational space as water molecules along the water array could both re-orient and undergo drying/wetting transitions. While conformational sampling with US was locked into pre-defined pathways, the MWE approach allowed us to map both direct- and water-mediated reaction pathways. This enhancement could provide a benefit in modeling different reaction mechanisms in complex systems, although it also leads to a significantly increased computational cost.

We also found that the system can be biased along the mCEC reaction coordinate such that the sampling displaces intervening water molecules from the “Grotthuss chain”. As a consequence, barriers orthogonal to the reaction coordinate are more easily overcome by MWE as compared to US. By introducing positional restraints on the groups participating in the proton transfer reactions, we could show that the US PMF is consistent with the SWE PMF (Figure S14). Indeed, the QM/MM-US method requires also pre-knowledge of suitable biasing potentials to achieve uniform sampling of the free energy landscape. In this regard, improved initial guesses of the optimal biasing potentials, *e*.*g*., from a MWE simulation could significantly enhance the rather slow exploration of the reaction in the QM/MM-US simulations.

To address the sampling and convergence issues, we propose a hybrid MWE/focused US approach for exploration of complex reaction mechanisms. To this end, QM/MM-MWE are performed to explore various putative reaction mechanisms. After identifying different pathways, the PMF is converged using QM/MM-US, whilst the PMF of the combined data set is obtained using MBAR or related techniques. The high degree of conformational changes and hydration variability in biological systems as observed in atomistic MD simulations highlights the necessity of exploring a wide region of conformational space during free energy sampling and provides a significant challenge particularly for DFT-based QM/MM-free energy calculations.

In contrast to previous approaches in which, *e*.*g*., metadynamics-explored pathways are subsequently sampled with US,^75, 76^ the current scheme developed here enables the incorporation of information gained from both MWE and FUS. This combination has the benefit of providing good initial estimates of the PMF with MWE, which can then be greatly improved in the FUS step. Usage of computational resources is thus enhanced, as both extensive and intensive sampling is performed. Nevertheless, FUS also requires a detailed understanding of the studied system in conjunction with a clearly defined target region of the reaction coordinate. Since the MBAR is employed as reweighting procedure, an arbitrary number of US windows can be simulated, allowing for the selection of multiple target regions.

In addition to providing barriers and driving forces, data obtained from free energy calculations can be used to describe the structures of relevant intermediates along the reaction pathways. We suggest that reaction intermediates obtained from the MWE sampling can be extracted using clustering methods. However, the direct time-ordering of physically realistic intermediates for derivation of reaction mechanisms is perhaps more straightforward in the QM/MM-US simulations, as the biasing force reaches a local equilibrium in each simulation window, thus allowing for the characterization of reaction steps along the collective variable. The mCEC/MWE approach suggests that some of the protonation reactions in Complex I may occur *via* multiple competing reaction pathways, possibly modulated by the hydration and protonation states of the surrounding groups. Although this requires further detailed exploration, it suggests interesting gating principles that could be used to modulate proton conduction, *e*.*g*., during reversal of the proton pumping machinery, which takes place during hypoxic conditions in mitochondria.

## CONLUSIONS

We have introduced, implemented, and compared here the multiple walker/well-tempered metadynamics – extended adaptive biasing force (MWE) and umbrella sampling (US) methods for GPU-accelerated QM/MM free energy calculations of water-mediated proton transfer reactions in a complex (bio)chemical system. To this end, we studied the performance of both MWE and US in combination with a geometric and a physical description of the proton transfer reaction coordinate. Our combined findings show that the MWE approach combined with the modified center of excess charge (mCEC) reaction coordinate can efficiently sample the protonation dynamics and orthogonal hydration dynamics, while the US method effectively sampled conformations containing the hydration pattern of the starting conformation. In contrast, we found that the mCEC/MWE approach achieves a multi-pathway exploration of the reaction coordinate that, for the studied model systems, converges within a timescale similar to that of the US method. For the proton transfer reactions in Complex I, we observe that convergence of the MWE sampling is increased relative to the US method, while combination of both approaches (QM/MM-MWE/FUS) provide an improved exploration of the 2D-PMF profiles. Our study provides key insights into the application of first principles-based QM/MM free energy methods for mechanistic studies of protonation dynamics in highly intricate biochemical systems, and highlights new mechanistic insight into protonation dynamics in Complex I.

## ASSOCIATED CONTENT

### Supporting Information

The Supporting Information shows convergence of the PMF, methods testing, and reaction coordinate values of central states for the studied systems. The Supporting Information is available free of charge on the ACS Publications website.

## AUTHOR INFORMATION

### Author Contributions

All authors have given approval to the final version of the manuscript.

## ACKNOWLEDGMENT

This work was funded by the Knut and Alice Wallenberg Foundation (grant: 2019.0251 and WASPDDLS22:025, VRIK), the Swedish Research Council (VRIK), and the German Research Foundation (DFG, TRR235, “Emergence of Life” to VRIK and CO), and by the European Research Council under the European Union’s Horizon 2020 research and innovation program/Grant Agreement 715311 (VRIK). JCBD acknowledges the support of the Leopoldina Fellowship Program, German National Academy of Sciences Leopoldina, grant number LPDS 2021-08. VRIK acknowledges support from the German Research Foundation (DFG) via the Collaborative Research Centre (SFB1078). This work was also supported by the National Academic Infrastructure for Supercomputing in Sweden (NAISS, NAISS: 2023/1-31, 2023/6-128) and the Swedish National Infrastructure for Computing (SNIC 2022/1-29, SNIC 2022/13-14) at the Center for High Performance Computing (PDC) Center, partially funded by the Swedish Research Council through grant agreements no. 2022-06725 and no. 2018-05973, and the Leibniz Rechenzentrum (LRZ, project:pr83ro), Germany.

## ABBREVIATIONS

PMF: potential of mean force
CV: collective variable
RC: reaction coordinate
LC: linear combination of bond-breaking and bond-formation
mCEC: modified center of excess charge
US: umbrella sampling
WTM-eABF: well-tempered metadynamics/extended adaptive biasing force
MWE: multiple-walkers/well-tempered metadynamics/extended adaptive biasing force
SWE: single-walker/well-tempered metadynamics/extended adaptive biasing force
QM/MM: hybrid quantum mechanics/molecular mechanics
MD: molecular dynamics
DFTB: density functional based tight binding
SCC-DFTB: self-consistent charge
DFTB, PM7: parametric method 7
EVB: empirical valence bond
MS-EVB: multi-state EVB.

---

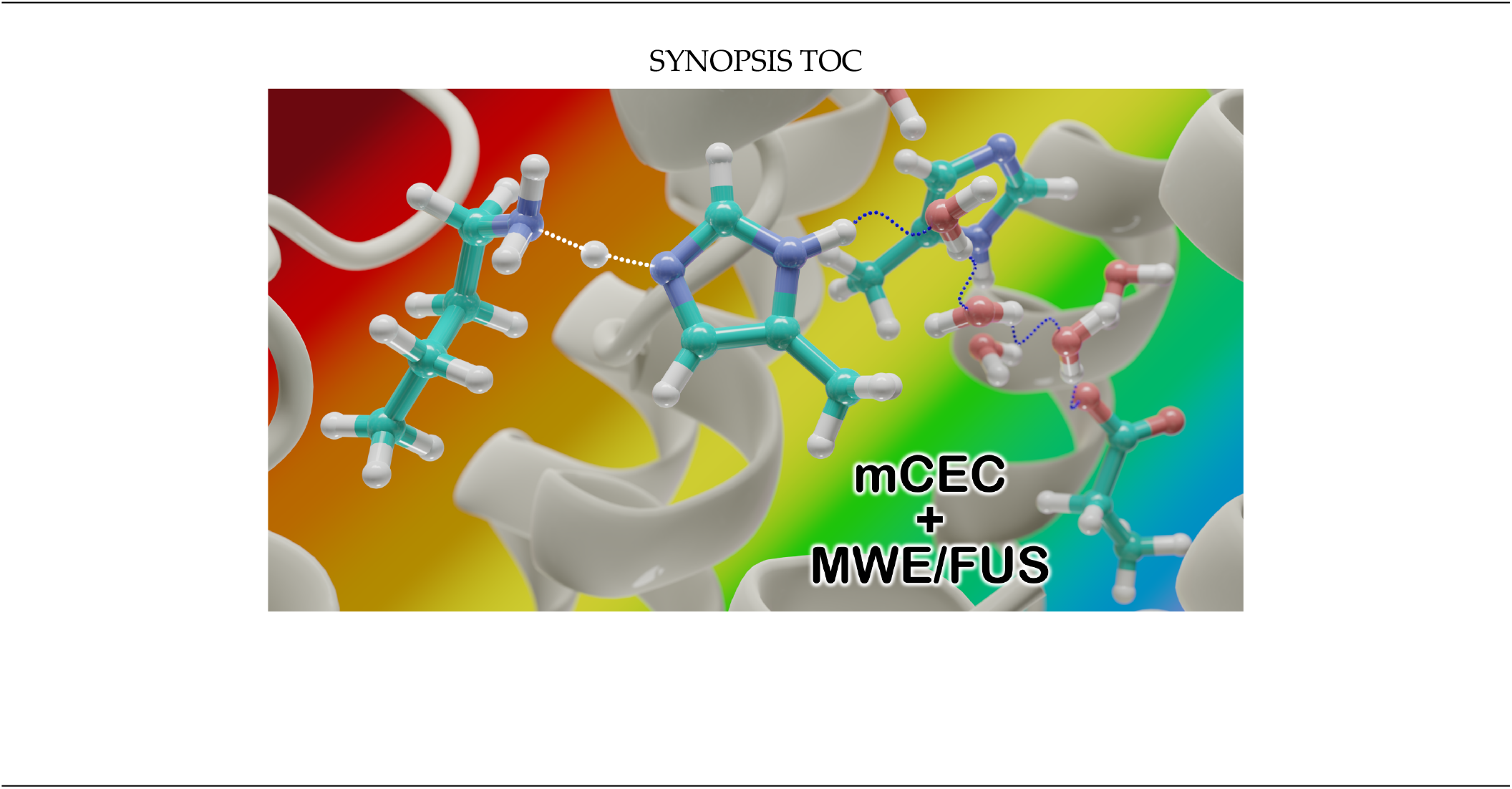

## REFERENCES

(1) Kaila, V. R. I. Resolving Chemical Dynamics in Biological Energy Conversion: Long-Range Proton-Coupled Electron Transfer in Respiratory Complex I. Accounts of Chemical Research 2021, 54 (24), 4462–4473. DOI: 10.1021/acs.accounts.1c00524.

(2) Kaila, V. R. I.; Wikström, M. Architecture of bacterial respiratory chains. Nature Reviews Microbiology 2021, 19 (5), 319–330. DOI: 10.1038/s41579-020-00486-4.

(3) Agmon, N. The Grotthuss mechanism. Chemical Physics Letters 1995, 244 (5), 456–462. DOI: 10.1016/0009-2614(95)00905-J.

(4) Mitchell, P. Coupling of Phosphorylation to Electron and Hydrogen Transfer by a Chemi-Osmotic type of Mechanism. Nature 1961, 191 (4784), 144–148. DOI: 10.1038/191144a0.

(5) Röpke, M.; Saura, P.; Riepl, D.; Pöverlein, M. C.; Kaila, V. R. I. Functional Water Wires Catalyze Long-Range Proton Pumping in the Mammalian Respiratory Complex I. J. Am. Chem. Soc. 2020, 142 (52), 21758–21766. DOI: 10.1021/jacs.0c09209.

(6) Röpke, M.; Riepl, D.; Saura, P.; Di Luca, A.; Mühlbauer, M. E.; Jussupow, A.; Gamiz-Hernandez, A. P.; Kaila, V. R. I. Deactivation blocks proton pathways in the mitochondrial complex I. Proceedings of the National Academy of Sciences 2021, 118 (29), e2019498118. DOI: 10.1073/pnas.2019498118.

(7) Mader, S. L.; Lopez, A.; Lawatscheck, J.; Luo, Q.; Rutz, D. A.; Gamiz-Hernandez, A. P.; Sattler, M.; Buchner, J.; Kaila, V. R. I. Conformational dynamics modulate the catalytic activity of the molecular chaperone Hsp90. Nature Communications 2020, 11 (1), 1410. DOI: 10.1038/s41467-020-15050-0.

(8) Liang, R.; Swanson, J. M. J.; Peng, Y.; Wikström, M.; Voth, G. A. Multiscale simulations reveal key features of the proton-pumping mechanism in cytochrome c oxidase. Proceedings of the National Academy of Sciences 2016, 113 (27), 7420–7425. DOI: 10.1073/pnas.1601982113.

(9) Li, C.; Yue, Z.; Espinoza-Fonseca, L. M.; Voth, G. A. Multiscale Simulation Reveals Passive Proton Transport Through SERCA on the Microsecond Timescale. Biophysical Journal 2020, 119 (5), 1033–1040. DOI: 10.1016/j.bpj.2020.07.027.

(10) Pisliakov, A. V.; Sharma, P. K.; Chu, Z. T.; Haranczyk, M.; Warshel, A. Electrostatic basis for the unidirectionality of the primary proton transfer in cytochrome c oxidase. Proceedings of the National Academy of Sciences 2008, 105 (22), 7726–7731. DOI: 10.1073/pnas.0800580105.

(11) Maag, D.; Mast, T.; Elstner, M.; Cui, Q.; Kubař, T. O to bR transition in bacteriorhodopsin occurs through a proton hole mechanism. Proceedings of the National Academy of Sciences 2021, 118 (39), e2024803118. DOI: 10.1073/pnas.2024803118.

(12) Elstner, M. The SCC-DFTB method and its application to biological systems. Theoretical Chemistry Accounts 2006, 116 (1), 316–325. DOI: 10.1007/s00214-005-0066-0.

(13) Mlýnský, V.; Banáš, P.; Šponer, J.; van der Kamp, M. W.; Mulholland, A. J.; Otyepka, M. Comparison of ab Initio, DFT, and Semiempirical QM/MM Approaches for Description of Catalytic Mechanism of Hairpin Ribozyme. Journal of Chemical Theory and Computation 2014, 10 (4), 1608–1622. DOI: 10.1021/ct401015e.

(14) Elstner, M.; Porezag, D.; Jungnickel, G.; Elsner, J.; Haugk, M.; Frauenheim, T.; Suhai, S.; Seifert, G. Self-consistent-charge density-functional tight-binding method for simulations of complex materials properties. Physical Review B 1998, 58 (11), 7260–7268. DOI: 10.1103/PhysRevB.58.7260.

(15) Stewart, J. J. P. Optimization of parameters for semiempirical methods VI: more modifications to the NDDO approximations and re-optimization of parameters. Journal of Molecular Modeling 2013, 19 (1), 1–32. DOI: 10.1007/s00894-012-1667-x.

(16) Christensen, A. S.; Kubař, T.; Cui, Q.; Elstner, M. Semiempirical Quantum Mechanical Methods for Noncovalent Interactions for Chemical and Biochemical Applications. Chemical Reviews 2016, 116 (9), 5301–5337. DOI: 10.1021/acs.chemrev.5b00584.

(17) Cohen, A. J.; Mori-Sánchez, P.; Yang, W. Challenges for Density Functional Theory. Chemical Reviews 2012, 112 (1), 289–320. DOI: 10.1021/cr200107z.

(18) Siegbahn, P. E. M.; Blomberg, M. R. A. Transition-Metal Systems in Biochemistry Studied by High-Accuracy Quantum Chemical Methods. Chemical Reviews 2000, 100 (2), 421–438. DOI: 10.1021/cr980390w.

(19) Kubař, T.; Elstner, M.; Cui, Q. Hybrid Quantum Mechanical/Molecular Mechanical Methods For Studying Energy Transduction in Biomolecular Machines. Annual Review of Biophysics 2023, 52 (1), 525–551. DOI: 10.1146/annurev-biophys-111622-091140.

(20) Kussmann, J.; Ochsenfeld, C. Preselective Screening for Linear-Scaling Exact Exchange-Gradient Calculations for Graphics Processing Units and General Strong-Scaling Massively Parallel Calculations. Journal of Chemical Theory and Computation 2015, 11 (3), 918–922. DOI: 10.1021/ct501189u.

(21) Kussmann, J.; Ochsenfeld, C. Hybrid CPU/GPU Integral Engine for Strong-Scaling Ab Initio Methods. Journal of Chemical Theory and Computation 2017, 13 (7), 3153–3159. DOI: 10.1021/acs.jctc.6b01166.

(22) Fu, H.; Shao, X.; Cai, W.; Chipot, C. Taming Rugged Free Energy Landscapes Using an Average Force. Accounts of Chemical Research 2019, 52 (11), 3254–3264. DOI: 10.1021/acs.accounts.9b00473.

(23) König, P. H.; Ghosh, N.; Hoffmann, M.; Elstner, M.; Tajkhorshid, E.; Frauenheim, T.; Cui, Q. Toward Theoretical Analyis of Long-Range Proton Transfer Kinetics in Biomolecular Pumps. The Journal of Physical Chemistry A 2006, 110 (2), 548–563. DOI: 10.1021/jp052328q.

(24) Laio, A.; Parrinello, M. Escaping free-energy minima. Proceedings of the National Academy of Sciences 2002, 99 (20), 12562–12566. DOI: 10.1073/pnas.202427399.

(25) Darve, E.; Pohorille, A. Calculating free energies using average force. The Journal of Chemical Physics 2001, 115 (20), 9169–9183. DOI: 10.1063/1.1410978.

(26) Sørensen, M. R.; Voter, A. F. Temperature-accelerated dynamics for simulation of infrequent events. The Journal of Chemical Physics 2000, 112 (21), 9599–9606. DOI: 10.1063/1.481576.

(27) Torrie, G. M.; Valleau, J. P. Nonphysical sampling distributions in Monte Carlo free-energy estimation: Umbrella sampling. Journal of Computational Physics 1977, 23 (2), 187–199. DOI: 10.1016/0021-9991(77)90121-8.

(28) Lesage, A.; Lelièvre, T.; Stoltz, G.; Hénin, J. Smoothed Biasing Forces Yield Unbiased Free Energies with the Extended-System Adaptive Biasing Force Method. The Journal of Physical Chemistry B 2017, 121 (15), 3676–3685. DOI: 10.1021/acs.jpcb.6b10055.

(29) Hulm, A.; Dietschreit, J. C. B.; Ochsenfeld, C. Statistically optimal analysis of the extended-system adaptive biasing force (eABF) method. The Journal of Chemical Physics 2022, 157 (2). DOI: 10.1063/5.0095554.

(30) Shirts, M. R.; Chodera, J. D. Statistically optimal analysis of samples from multiple equilibrium states. The Journal of Chemical Physics 2008, 129 (12). DOI: 10.1063/1.2978177.

(31) Wu, H.; Paul, F.; Wehmeyer, C.; Noé, F. Multiensemble Markov models of molecular thermodynamics and kinetics. Proceedings of the National Academy of Sciences 2016, 113 (23), E3221–E3230. DOI: 10.1073/pnas.1525092113.

(32) Wu, H.; Mey, A. S. J. S.; Rosta, E.; Noé, F. Statistically optimal analysis of state-discretized trajectory data from multiple thermodynamic states. The Journal of Chemical Physics 2014, 141 (21). DOI: 10.1063/1.4902240.

(33) Rosta, E.; Hummer, G. Free Energies from Dynamic Weighted Histogram Analysis Using Unbiased Markov State Model. Journal of Chemical Theory and Computation 2015, 11 (1), 276–285. DOI: 10.1021/ct500719p.

(34) Stelzl, L. S.; Kells, A.; Rosta, E.; Hummer, G. Dynamic Histogram Analysis To Determine Free Energies and Rates from Biased Simulations. Journal of Chemical Theory and Computation 2017, 13 (12), 6328–6342. DOI: 10.1021/acs.jctc.7b00373.

(35) Dietschreit, J. C. B.; Diestler, D. J.; Hulm, A.; Ochsenfeld, C.; Gómez-Bombarelli, R. From free-energy profiles to activation free energies. The Journal of Chemical Physics 2022, 157 (8). DOI: 10.1063/5.0102075.

(36) Becke, A. D. Density-functional exchange-energy approximation with correct asymptotic behavior. Physical Review A 1988, 38 (6), 3098–3100. DOI: 10.1103/PhysRevA.38.3098.

(37) Lee, C.; Yang, W.; Parr, R. G. Development of the Colle-Salvetti correlation-energy formula into a functional of the electron density. Phys. Rev. B 1988, 37 (2), 785–789. DOI: 10.1103/physrevb.37.785.

(38) Weigend, F.; Ahlrichs, R. Balanced basis sets of split valence, triple zeta valence and quadruple zeta valence quality for H to Rn: Design and assessment of accuracy. Physical Chemistry Chemical Physics 2005, 7 (18), 3297–3305. DOI: 10.1039/B508541A.

(39) Grimme, S.; Antony, J.; Ehrlich, S.; Krieg, H. A consistent and accurate ab initio parametrization of density functional dispersion correction (DFT-D) for the 94 elements H-Pu. J. Chem. Phys. 2010, 132 (15), 154104. DOI: 10.1063/1.3382344.

(40) Mangiatordi, G. F.; Brémond, E.; Adamo, C. DFT and Proton Transfer Reactions: A Benchmark Study on Structure and Kinetics. Journal of Chemical Theory and Computation 2012, 8 (9), 3082–3088. DOI: 10.1021/ct300338y.

(41) David, R.; Jamet, H.; Nivière, V.; Moreau, Y.; Milet, A. Iron Hydroperoxide Intermediate in Superoxide Reductase: Protonation or Dissociation First? MM Dynamics and QM/MM Metadynamics Study. Journal of Chemical Theory and Computation 2017, 13 (6), 2987–3004. DOI: 10.1021/acs.jctc.7b00126.

(42) Duster, A. W.; Lin, H. Tracking Proton Transfer through Titratable Amino Acid Side Chains in Adaptive QM/MM Simulations. Journal of Chemical Theory and Computation 2019, 15 (11), 5794–5809. DOI: 10.1021/acs.jctc.9b00649.

(43) Mühlbauer, M. E.; Saura, P.; Nuber, F.; Di Luca, A.; Friedrich, T.; Kaila, V. R. I. Water-Gated Proton Transfer Dynamics in Respiratory Complex I. Journal of the American Chemical Society 2020, 142 (32), 13718–13728. DOI: 10.1021/jacs.0c02789.

(44) Yagi, K.; Ito, S.; Sugita, Y. Exploring the Minimum-Energy Pathways and Free-Energy Profiles of Enzymatic Reactions with QM/MM Calculations. The Journal of Physical Chemistry B 2021, 125 (18), 4701–4713. DOI: 10.1021/acs.jpcb.1c01862.

(45) Dürr, S. L.; Bohuszewicz, O.; Berta, D.; Suardiaz, R.; Jambrina, P. G.; Peter, C.; Shao, Y.; Rosta, E. The Role of Conserved Residues in the DEDDh Motif: the Proton-Transfer Mechanism of HIV-1 RNase H. ACS Catalysis 2021, 11 (13), 7915–7927. DOI: 10.1021/acscatal.1c01493.

(46) Kim, H.; Saura, P.; Pöverlein, M. C.; Gamiz-Hernandez, A. P.; Kaila, V. R. I. Quinone Catalysis Modulates Proton Transfer Reactions in the Membrane Domain of Respiratory Complex I. Journal of the American Chemical Society 2023, 145 (31), 17075–17086. DOI: 10.1021/jacs.3c03086.

(47) Sheng, X.; Himo, F. The Quantum Chemical Cluster Approach in Biocatalysis. Accounts of Chemical Research 2023, 56 (8), 938–947. DOI: 10.1021/acs.accounts.2c00795.

(48) Siegbahn, P. E. M. A quantum chemical approach for the mechanisms of redox-active metalloenzymes. RSC Advances 2021, 11 (6), 3495-3508, 10.1039/D0RA10412D. DOI: 10.1039/D0RA10412D.

(49) Klamt, A.; Schüürmann, G. COSMO: a new approach to dielectric screening in solvents with explicit expressions for the screening energy and its gradient. Journal of the Chemical Society, Perkin Transactions 2 1993, (5), 799–805. DOI: 10.1039/P29930000799.

(50) Shirts, M. R.; Ferguson, A. L. Statistically Optimal Continuous Free Energy Surfaces from Biased Simulations and Multistate Reweighting. Journal of Chemical Theory and Computation 2020, 16 (7), 4107–4125. DOI: 10.1021/acs.jctc.0c00077.

(51) Kussmann, J.; Ochsenfeld, C. Pre-selective screening for matrix elements in linear-scaling exact exchange calculations. The Journal of Chemical Physics 2013, 138 (13). DOI: 10.1063/1.4796441.

(52) Laqua, H.; Kussmann, J.; Ochsenfeld, C. Accelerating seminumerical Fock-exchange calculations using mixed single- and double-precision arithmethic. The Journal of Chemical Physics 2021, 154 (21). DOI: 10.1063/5.0045084.

(53) Kussmann, J.; Laqua, H.; Ochsenfeld, C. Highly Efficient Resolution-of-Identity Density Functional Theory Calculations on Central and Graphics Processing Units. Journal of Chemical Theory and Computation 2021, 17 (3), 1512–1521. DOI: 10.1021/acs.jctc.0c01252.

(54) Eastman, P.; Swails, J.; Chodera, J. D.; McGibbon, R. T.; Zhao, Y.; Beauchamp, K. A.; Wang, L.-P.; Simmonett, A. C.; Harrigan, M. P.; Stern, C. D.; et al. OpenMM 7: Rapid development of high performance algorithms for molecular dynamics. PLOS Computational Biology 2017, 13 (7), e1005659. DOI: 10.1371/journal.pcbi.1005659.

(55) Laqua, H.; Thompson, T. H.; Kussmann, J.; Ochsenfeld, C. Highly Efficient, Linear-Scaling Seminumerical Exact-Exchange Method for Graphic Processing Units. Journal of Chemical Theory and Computation 2020, 16 (3), 1456–1468. DOI: 10.1021/acs.jctc.9b00860.

(56) Laqua, H.; Dietschreit, J. C. B.; Kussmann, J.; Ochsenfeld, C. Accelerating Hybrid Density Functional Theory Molecular Dynamics Simulations by Seminumerical Integration, Resolution-of-the-Identity Approximation, and Graphics Processing Units. Journal of Chemical Theory and Computation 2022, 18 (10), 6010–6020. DOI: 10.1021/acs.jctc.2c00509.

(57) TURBOMOLE V7.4 2019, a development of University of Karlsruhe and Forschungszentrum Karlsruhe GmbH, 1989-2007, TURBOMOLE GmbH, since 2007; available from http://www.turbomole.com.

(58) Ahlrichs, R.; Bär, M.; Häser, M.; Horn, H.; Kölmel, C. Electronic structure calculations on workstation computers: The program system turbomole. Chemical Physics Letters 1989, 162 (3), 165–169. DOI: 10.1016/0009-2614(89)85118-8.

(59) Humphrey, W.; Dalke, A.; Schulten, K. VMD: Visual molecular dynamics. Journal of Molecular Graphics 1996, 14 (1), 33–38. DOI: 10.1016/0263-7855(96)00018-5.

(60) Delano, W. L. PyMOL Molecular Graphics System . Schrödinger LLC, 2002. https://sourceforge.net/projects/pymol/.

(61) Bannwarth, C.; Ehlert, S.; Grimme, S. GFN2-xTB—An Accurate and Broadly Parametrized Self-Consistent Tight-Binding Quantum Chemical Method with Multipole Electrostatics and Density-Dependent Dispersion Contributions. Journal of Chemical Theory and Computation 2019, 15 (3), 1652–1671. DOI: 10.1021/acs.jctc.8b01176.

(62) Ehlert, S.; Stahn, M.; Spicher, S.; Grimme, S. Robust and Efficient Implicit Solvation Model for Fast Semiempirical Methods. Journal of Chemical Theory and Computation 2021, 17 (7), 4250–4261. DOI: 10.1021/acs.jctc.1c00471.

(63) Bridges, H. R.; Fedor, J. G.; Blaza, J. N.; Di Luca, A.; Jussupow, A.; Jarman, O. D.; Wright, J. J.; Agip, A.-N. A.; Gamiz-Hernandez, A. P.; Roessler, M. M.; et al. Structure of inhibitor-bound mammalian complex I. Nature Communications 2020, 11 (1), 5261. DOI: 10.1038/s41467-020-18950-3.

(64) Best, R. B.; Zhu, X.; Shim, J.; Lopes, P. E. M.; Mittal, J.; Feig, M.; MacKerell, A. D., Jr. Optimization of the Additive CHARMM All-Atom Protein Force Field Targeting Improved Sampling of the Backbone ϕ,? and Side-Chain χ1 and χ2 Dihedral Angles. J. Chem. Comput. Theo. 2012, 8 (9), 3257–3273. DOI: 10.1021/ct300400x.

(65) Pracht, P.; Bohle, F.; Grimme, S. Automated exploration of the low-energy chemical space with fast quantum chemical methods. Physical Chemistry Chemical Physics 2020, 22 (14), 7169–7192. DOI: 10.1039/C9CP06869D.

(66) Kaila, V. R. I.; Hummer, G. Energetics and dynamics of proton transfer reactions along short water wires. Physical Chemistry Chemical Physics 2011, 13 (29), 13207-13215, 10.1039/C1CP21112A. DOI: 10.1039/C1CP21112A.

(67) Kaila, V. R. I.; Hummer, G. Energetics of Direct and Water-Mediated Proton-Coupled Electron Transfer. Journal of the American Chemical Society 2011, 133 (47), 19040–19043. DOI: 10.1021/ja2082262.

(68) Saura, P.; Frey, D. M.; Gamiz-Hernandez, A. P.; Kaila, V. R. I. Electric field modulated redox-driven protonation and hydration energetics in energy converting enzymes. Chemical Communications 2019, 55 (43), 6078–6081. DOI: 10.1039/C9CC01135H.

(69) Berezhkovskii, A. M.; Szabo, A. Committors, first-passage times, fluxes, Markov states, milestones, and all that. The Journal of Chemical Physics 2019, 150 (5). DOI: 10.1063/1.5079742.

(70) Chupeau, M.; Gladrow, J.; Chepelianskii, A.; Keyser, U. F.; Trizac, E. Optimizing Brownian escape rates by potential shaping. Proceedings of the National Academy of Sciences 2020, 117 (3), 1383–1388. DOI: 10.1073/pnas.1910677116.

(71) Dietschreit, J. C. B.; Diestler, D. J.; Gómez-Bombarelli, R. Entropy and Energy Profiles of Chemical Reactions. Journal of Chemical Theory and Computation 2023, 19 (16), 5369–5379. DOI: 10.1021/acs.jctc.3c00448.

(72) Kaila, V. R. I. Long-range proton-coupled electron transfer in biological energy conversion: towards mechanistic understanding of respiratory complex I. Journal of The Royal Society Interface 2018, 15 (141), 20170916. DOI: 10.1098/rsif.2017.0916.

(73) Chung, I.; Grba, D. N.; Wright, J. J.; Hirst, J. Making the leap from structure to mechanism: are the open states of mammalian complex I identified by cryoEM resting states or catalytic intermediates? Current Opinion in Structural Biology 2022, 77, 102447. DOI: 10.1016/j.sbi.2022.102447.

(74) Kampjut, D.; Sazanov, L. A. Structure of respiratory complex I – An emerging blueprint for the mechanism. Current Opinion in Structural Biology 2022, 74, 102350. DOI: 10.1016/j.sbi.2022.102350.

(75) Autieri, E.; Sega, M.; Pederiva, F.; Guella, G. Puckering free energy of pyranoses: A NMR and metadynamics-umbrella sampling investigation. The Journal of Chemical Physics 2010, 133 (9). DOI: 10.1063/1.3476466.

(76) Babin, V.; Roland, C.; Darden, T. A.; Sagui, C. The free energy landscape of small peptides as obtained from metadynamics with umbrella sampling corrections. The Journal of Chemical Physics 2006, 125 (20). DOI: 10.1063/1.2393236.

