## Supplementary Information for "QM/MM Free Energy Calculations of Long-Range Biological Protonation Dynamics by Adaptive and Focused Sampling"


### Content

#### Extended Methods


Essential Dynamics Analysis (EDA)
Semi-Empirical Conformer Search
CV Mass and de Broglie thermal wavelength
Derivation of Eqn. 7 for the activation free energy for a 1D harmonic oscillator


#### Supplementary Figures


Figure S1 | Sampling the His-water model along the LC-CV.
Figure S2 | Sampling the His-water model along the mCEC-CV.
Figure S3 | Conformational sampling of proton transfer in the His-water model.
Figure S4 | Projection of the mCEC-CV on the LC-CV.
Figure S5 | EDA of the His-water model.
Figure S6 | Effect of switching parameter on the mCEC-CV and the PMF.
Figure S7 | Sampling of the proton transfer reaction in the membrane domain of Complex I with the US and MWE methods.
Figure S8 | Sampling of protonation states in the membrane domain of the Complex I model with the US and MWE methods.
Figure S9 | Intermediate structures sampled during proton transfer in Complex I.
Figure S10 | EDA of proton transfer reactions sampled in the membrane domain of Complex I.
Figure S11 | Conformer-rotamer search obtained in CREST sampling for the Complex I model.
Figure S12 | Sampled hydration states during proton transfer in the Complex I model.
Figure S13 | Sampled protonation states during proton transfer in the Complex I model.
Figure S14 | Sampling of proton transfer reactions in the Complex I model, with smaller QM region and positional restraints.
Figure S15 | Benchmarking the proton transfer energetics for the His-water model (model 1).


#### Supplementary Tables


Table S1 | List of simulations.
Table S2 | Apparent barriers and activation free energies for Model 1.


#### SI References

### Extended Methods

#### *Essential Dynamics Analysis*

Essential dynamics analysis (EDA) for the His-water model (Model 1; Figure 1) was performed on the Cartesian coordinates of all atoms in the system. For the proton transfer reaction in Complex I (Model 2), the EDA was performed on the Cartesian coordinates of the QM atoms of the QM/MM system (Figure 3). The EDA was performed using ProDy.[1] Similar to principal component analysis (PCA), the EDA orthogonalizes the covariance matrix to obtain principle modes of motion. The eigenvectors corresponding to the largest eigenvalues are interpreted as the dominant modes of motion.[2]

#### *Semi-Empirical Conformer Search*

Based on the initial conformation obtained from free energy sampling of the proton transfer reactions in Complex I, a conformational search was performed using the conformer-rotamer ensemble sampling tool (CREST).[3] The conformational search was performed on the QM region and its immediate surroundings, by extending the QM region from 114 atoms to 216 atoms (Figure S7A). The sidechains of amino acids were modeled by replacing the $C_\alpha$ atoms by hydrogen atoms, which were fixed to the $C_\alpha$ positions in the reference structure. After geometry optimization at the GFN2-xTB level of theory,[4] the conformational search was also performed at the GFN2-xTB level.[4]

#### *CV mass and de Broglie thermal wavelength*

The inverse mass of the pseudo-particle associated with the reaction coordinate can be defined as (*cf.* also Ref. [5], Eqn. 27),

$$m_\xi^{-1} = \left(\vec{\nabla}_{\boldsymbol{x}}\xi\right)^{\mathrm{T}}\mathbf{M}^{-1}\left(\vec{\nabla}_{\boldsymbol{x}}\xi\right) = \sum_{i}^{3N} m_i \left(\frac{\partial\xi}{\partial x_i}\right)^2 \qquad (1)$$

where $\vec{\nabla}_{\boldsymbol{x}}\xi$ denotes the gradient of the reaction coordinate with respect to the nuclear coordinates and $\mathbf{M}$ is the $3N \times 3N$ diagonal matrix of atomic masses. The de Broglie thermal wavelength of the reaction coordinate-associated pseudo-particle (Eqn. 7, *cf.* also Ref. [5]) is defined as,

$$\lambda_\xi \equiv \frac{h}{\sqrt{2\pi m_\xi k_B T}} \qquad (2)$$

where $T$ is the temperature of the system.

**Derivation of Eqn. 7. for the activation free energy for a 1D harmonic oscillator**

Let us assume that the potential at the reactant minimum ($R_{\text{min}}$) and transition state (TS) can be approximated harmonically by,

$$H_{R_{min}} = U_{R_{min}} + \frac{1}{2}(kq^2 + \frac{p^2}{m}) \qquad H_{TS} = U_{TS}$$

where $q$ is the normal mode and $p$ the conjugate momentum and $U_\alpha$ the potential energy of configuration, with the imaginary mode for the transition state excluded from the Hamiltonian. The activation free energy is the difference of the transition state and reactant free energies,

$$\Delta F^\ddagger = F_{TS} - F_R = -\beta^{-1}\ln\frac{Q_{TS}}{Q_R} = -\beta^{-1}\ln\frac{Z_{TS}\,\Lambda_R}{\Lambda_{TS}\,Z_R}$$

Using the harmonic Hamiltonians, the analytical configuration integrals $Z_\alpha$ and product of thermal wavelengths $\Lambda_\alpha$ for reactant and transition state are,

$$Z_R = e^{-\beta U_R}\int_{-\infty}^{\infty} dq\, e^{-\frac{\beta k q^2}{2}} = e^{-\beta U_R}\sqrt{\frac{2\pi}{\beta k}} \qquad Z_{TS} = e^{-\beta U_{TS}}$$

$$\Lambda_R^{\ -1} = \frac{1}{h}\int_{-\infty}^{\infty} dp\, e^{-\frac{\beta p^2}{2m}} = \sqrt{\frac{2\pi m}{\beta h^2}} \qquad \Lambda_{TS}^{\ -1} = 1$$

$Z_R$ and $Z_{TS}$ differ by the dimension of one normal mode, and $\Lambda_R$ and $\Lambda_{TS}$ by exactly one thermal wavelength. Hence, the argument of the logarithm in the equation for the activation free energy retains the thermal wavelength,

$$\Delta F^\ddagger = -\beta^{-1}\ln\left(\frac{e^{-\beta U_{TS}}\ \sqrt{\beta h^2/2\pi m}}{e^{-\beta U_R}\sqrt{2\pi/\beta k}}\right)$$

Comparing the harmonic approximation with Eq. 7 yields the corresponding terms,

$$\frac{Z_{TS}}{Z_R} = \frac{e^{-\beta U_{TS}}}{e^{-\beta U_R}\sqrt{2\pi/\beta k}} \ \widehat{=}\ \frac{\rho(z^\ddagger)}{P(\mathrm{R})} \quad \text{and} \quad \frac{\Lambda_R}{\Lambda_{TS}} = \sqrt{\beta h^2/2\pi m} \ \widehat{=}\ \langle\lambda_\xi\rangle_{z^\ddagger}$$

### Supplementary Figures

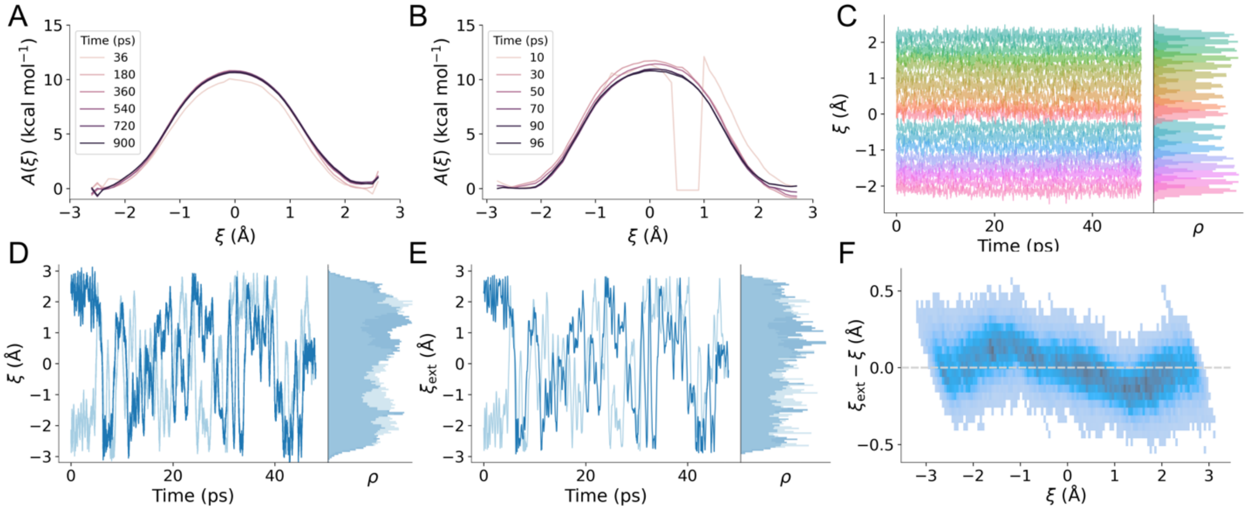


**Figure S1.** Sampling of the proton transfer reaction in the His-water model along the linear combination (LC) reaction coordinate. (A) Convergence of the PMF obtained from US. (B) Convergence of the PMF from MWE. (C) Sampling of the reaction coordinate using QM/MM-US. (D) Sampling of the reaction coordinate using QM/MM-MWE. (E) Sampling of the extended variable (eABF) reaction coordinate in MWE. (F) 2D histogram of the instantaneous difference between the extended system variable and the reaction coordinate as a function of the reaction coordinate.

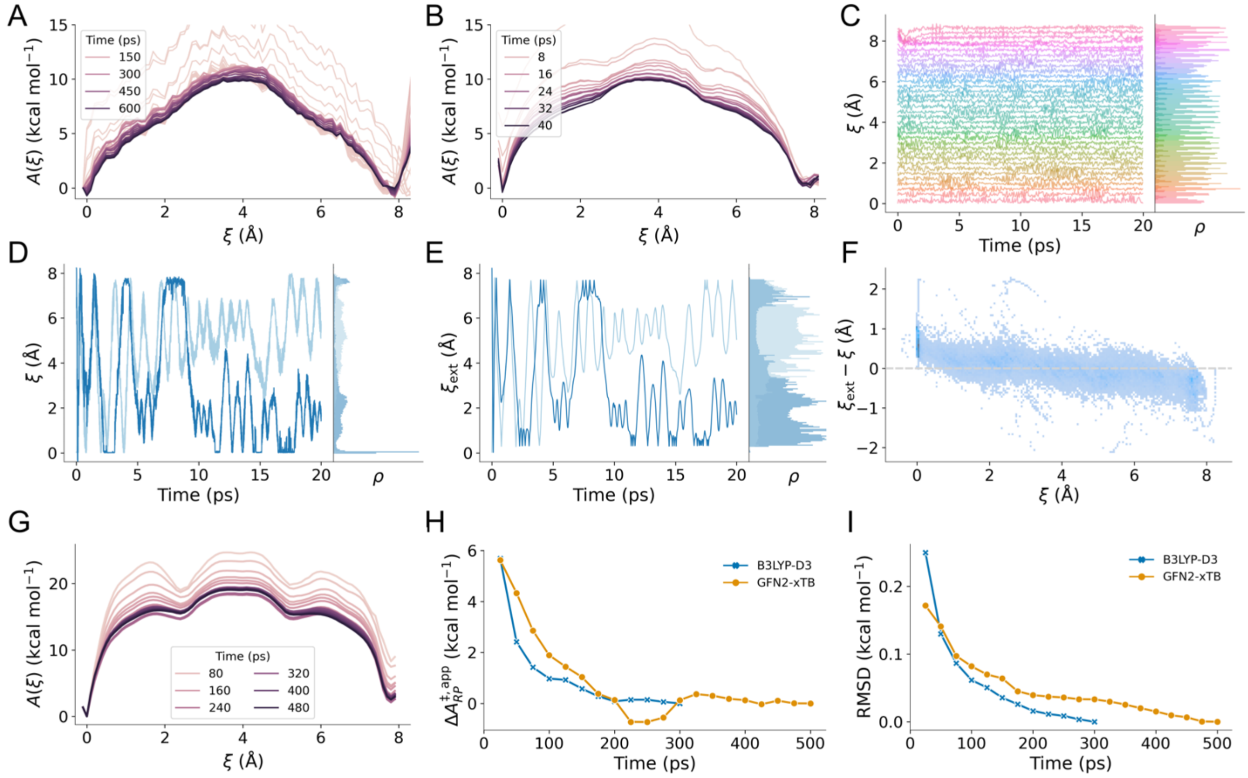


**Figure S2**. Sampling of the proton transfer reaction in the His-water model system using the modified center of excess charge reaction coordinate. (A) Convergence of the PMF obtained from US. (B) Convergence of the PMF from MWE. (C) Sampling of the extended variable reaction coordinate by US. (D) Sampling of the reaction coordinate using QM/MM-MWE. (E) Sampling of the extended variable (eABF) reaction coordinate in MWE. (F) 2D histogram of the instantaneous difference between the extended system variable and the reaction coordinate as a function of the reaction coordinate. (G) Convergence of the PMF from MWE at the GFN2-xTB level of theory. (H) Convergence of apparent barrier height ($\Delta A^{\ddagger,\mathrm{app}}_{RP} = A^{\ddagger} - A_R$) and (I) evolution of the PMF RMSD both with respect to final PMF profile.

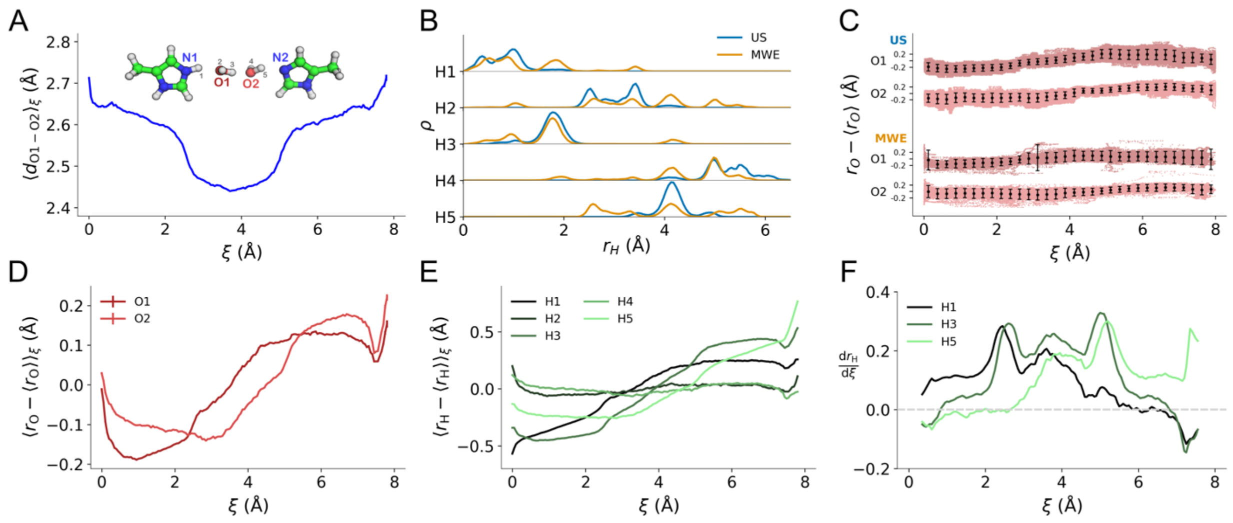


**Figure S3**. Conformational sampling of proton transfer in the His-water model. (A) Weighted average of O-O distance. (B) Histogram of the proton positions projected onto the donor-acceptor vector. (C) Projection of the water-O1 and water-O2 positions onto the reaction coordinate vector plotted versus reaction coordinate value. Error bars indicate 5th and 95th percentile. (D) Relative position of the O atoms projected onto the donor-acceptor vector. (E) Projected relative positions of the H atoms. (F) Derivative of the projected positions of the H atoms H1, H3, and H5. MWE data shown in A-F were obtained from 300 ps run of MWE.

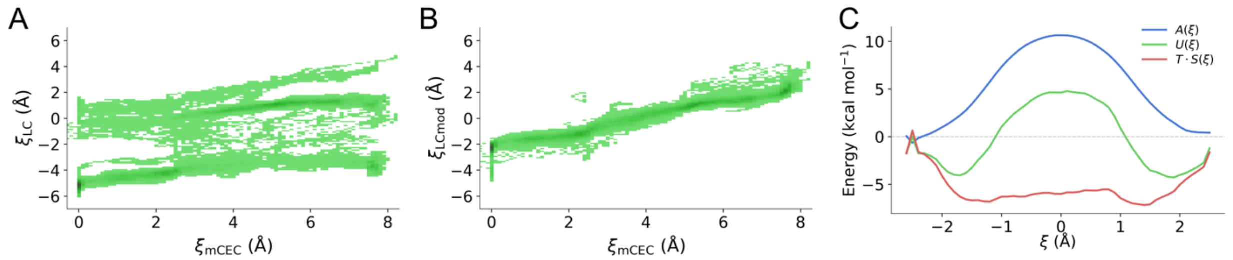


**Figure S4**. Projection of the mCEC-CV on the LC-CV. (A) Mapping of the mCEC sampling onto LC reaction coordinate. Reactant and product minima are indicated by dashed lines. Proton exchange during mCEC sampling leads to absence of separation between reactant and product state for the LC RC. (B) Projection of a modified LC reaction coordinate re-maps the reactant and product state to the physical RC range. (C) Free energy *A*, internal energy *U*, and entropy *T*·*S* profile of the His-water model as a function of the mCEC coordinate sampled with MWE.

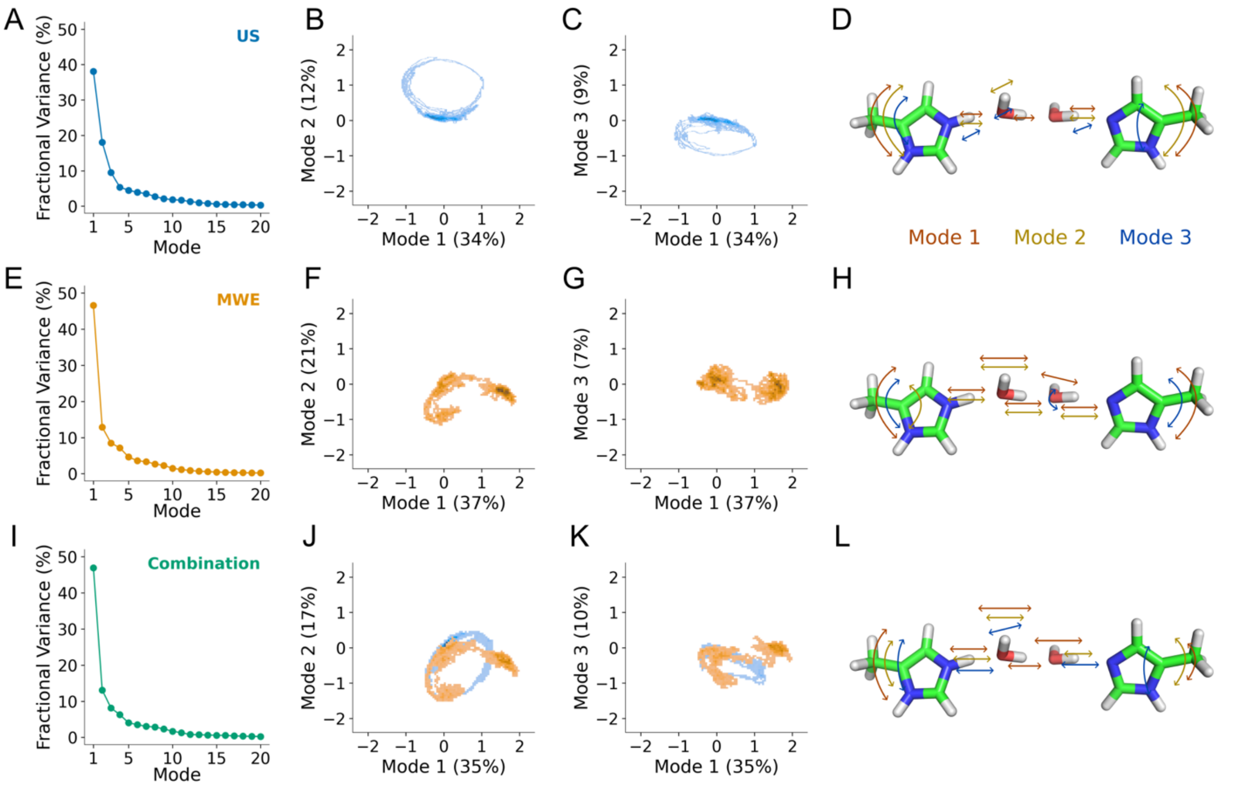


**Figure S5**. Essential dynamics analysis of the His-water model with the modified center of excess charge reaction coordinate using (A-D) US, (E-H) MWE. (I-L) Combination of the data from US and MWE. (A, E, I) Fractional variance of the first 20 principal components. (B, F, J) Projection of the trajectory data onto PC 1 and PC 2. (C, G, K) Projection of the trajectory data onto PC 1 and PC 3. (D, H, L) Visualization of the movements described by PC 1, PC 2, and PC 3.

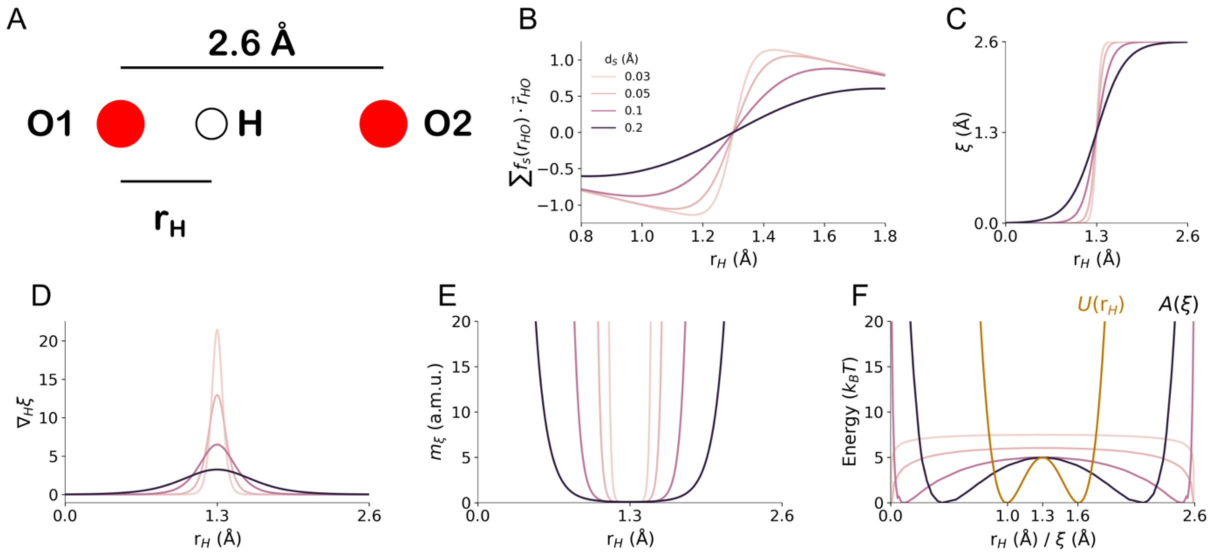


**Figure S6**. Effect of switching parameter on mCEC-CV and PMF. (A) Illustration of the of a O-H-O test system. (B) Sum of the mCEC correction terms as a function of the hydrogen position $r_H$ for different $d_s$. (C) mCEC-CV value as a function of $r_H$. (D) Gradient of the mCEC with respect to $r_H$. (E) Mass of the mCEC. (F) PMFs $A(\xi;d_s)$ when the model system is sampled assuming a double-well potential (red) $U(x)$=565.8436 ($x^4$ - 5.2$x^3$ + 9.9518$x^2$ - 8.2987$x$ + 2.5469), which was obtained by fitting a fourth degree polynomial to the points ((0.7,40), (1,0), (1.3,5), (1.6,0), (1.9,40)).

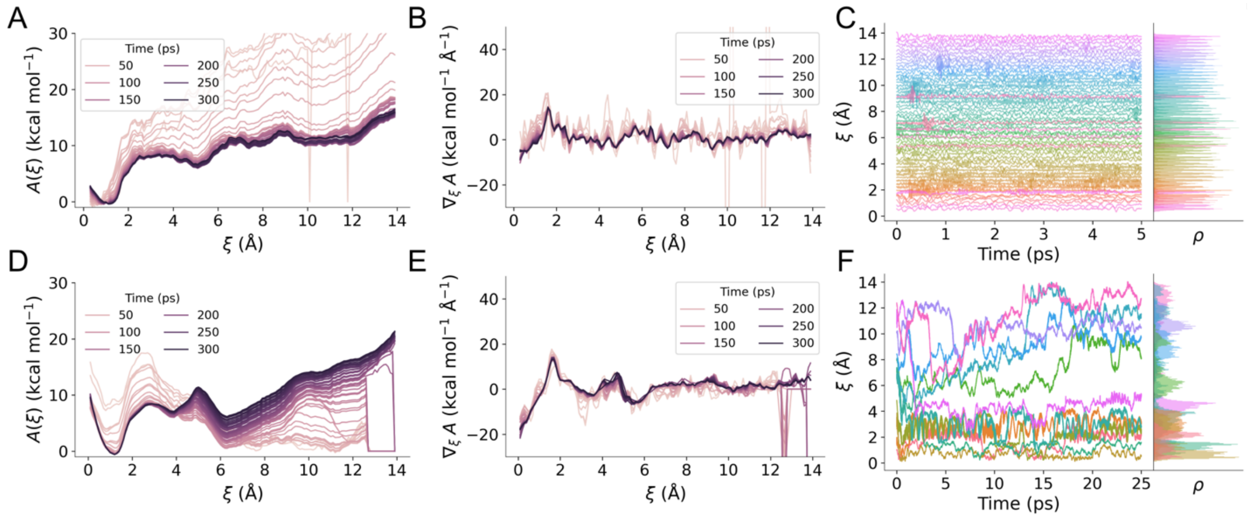


**Figure S7.** Sampling of the proton transfer reactions in the membrane domain of Complex I with the US and MWE methods. (A,D) Convergence of the PMF obtained from US and MWE, respectively. (B,E) Convergence of the mean force profile obtained from US and MWE, respectively. (C,F) Sampling of the reaction coordinate using US and MWE.

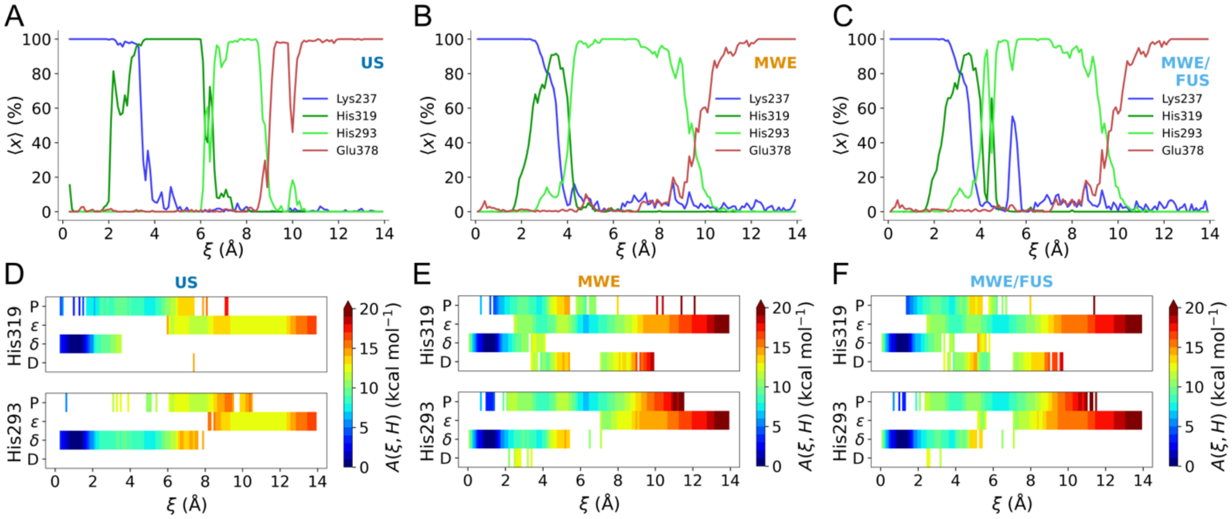


**Figure S8.** Sampling of protonation states in the membrane domain of Complex I with the US and MWE methods. (A,B,C) Protonation probability of the titratable residues as a function of the collective variable for US, MWE, and MWE/FUS, respectively. (D,E,F) Protonation state PMF profile of His319 (top) and His293 (bottom) for US, MWE, and MWE/FUS, respectively.

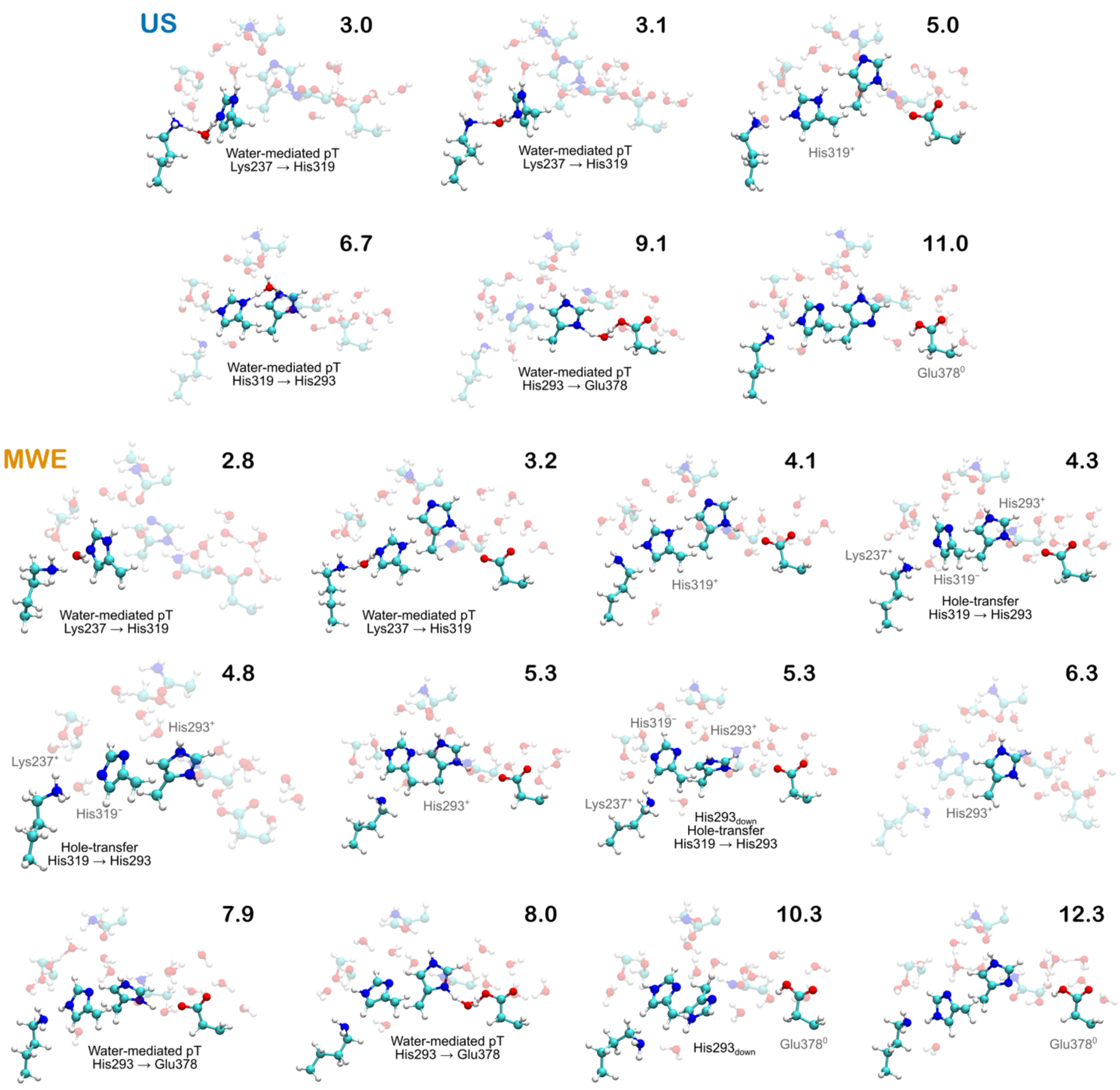


**Figure S9**. Intermediate structures sampled during proton transfer in Complex I from US and MWE. RC values (in Å) are indicated for each conformation.

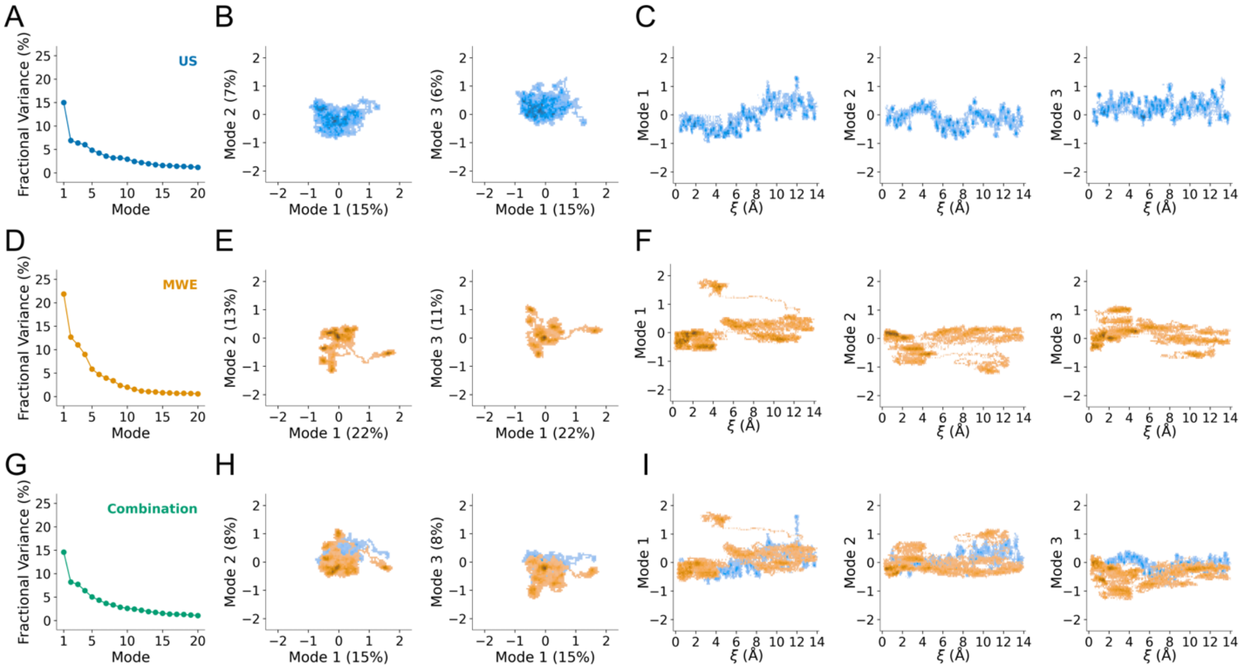


**Figure S10.** Essential dynamics analysis of proton transfer reactions sampled in the membrane domain of Complex I with the modified center of excess charge reaction coordinate using (A-C) US, (D-F) MWE, (G-I) combination of the data from US and MWE. (A, D, G) Fractional variances of principal component analysis of the QM region (B, E, H) Projection of the trajectory plotted onto principal components 1 and 2 (left) and 1 and 3 (right). (C, F, I) Projection of the trajectory onto principal components 1/2/3 as a function of collective variable.

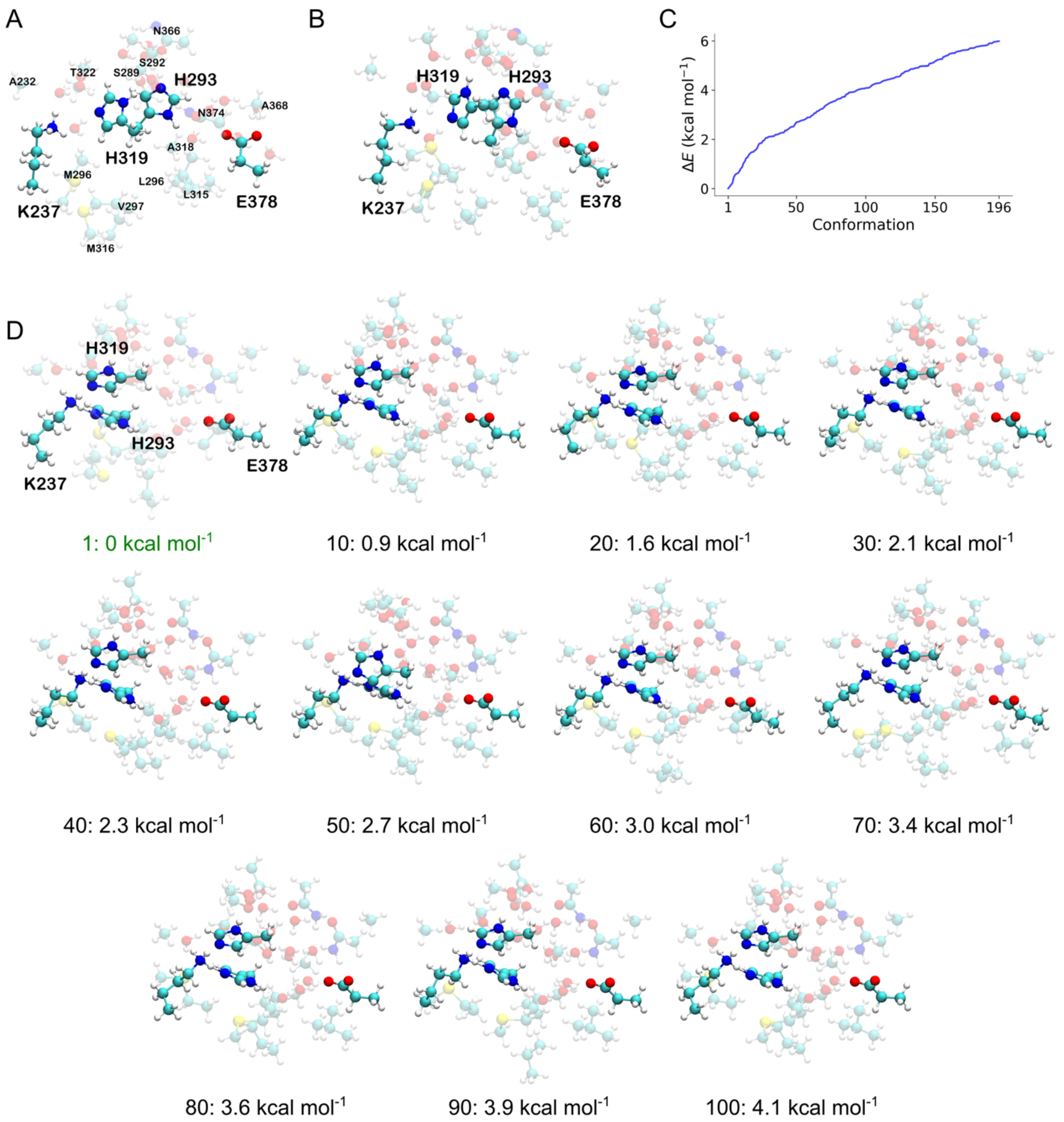


**Figure S11**. Conformer-rotamer search obtained in CREST sampling for the Complex I model. (A) Input conformation and residue numbering. (B) Structure obtained after optimization at the GFN2-xTB level. (C) Relative energies of the 196 conformers found below the threshold of 6 kcal $mol^{-1}$. (D) Every $10^{th}$ conformer from the initial 100 conformers, labeled with their respective energy differences in comparison to the lowest energy conformer. The lowest energy conformer is highlighted in green.

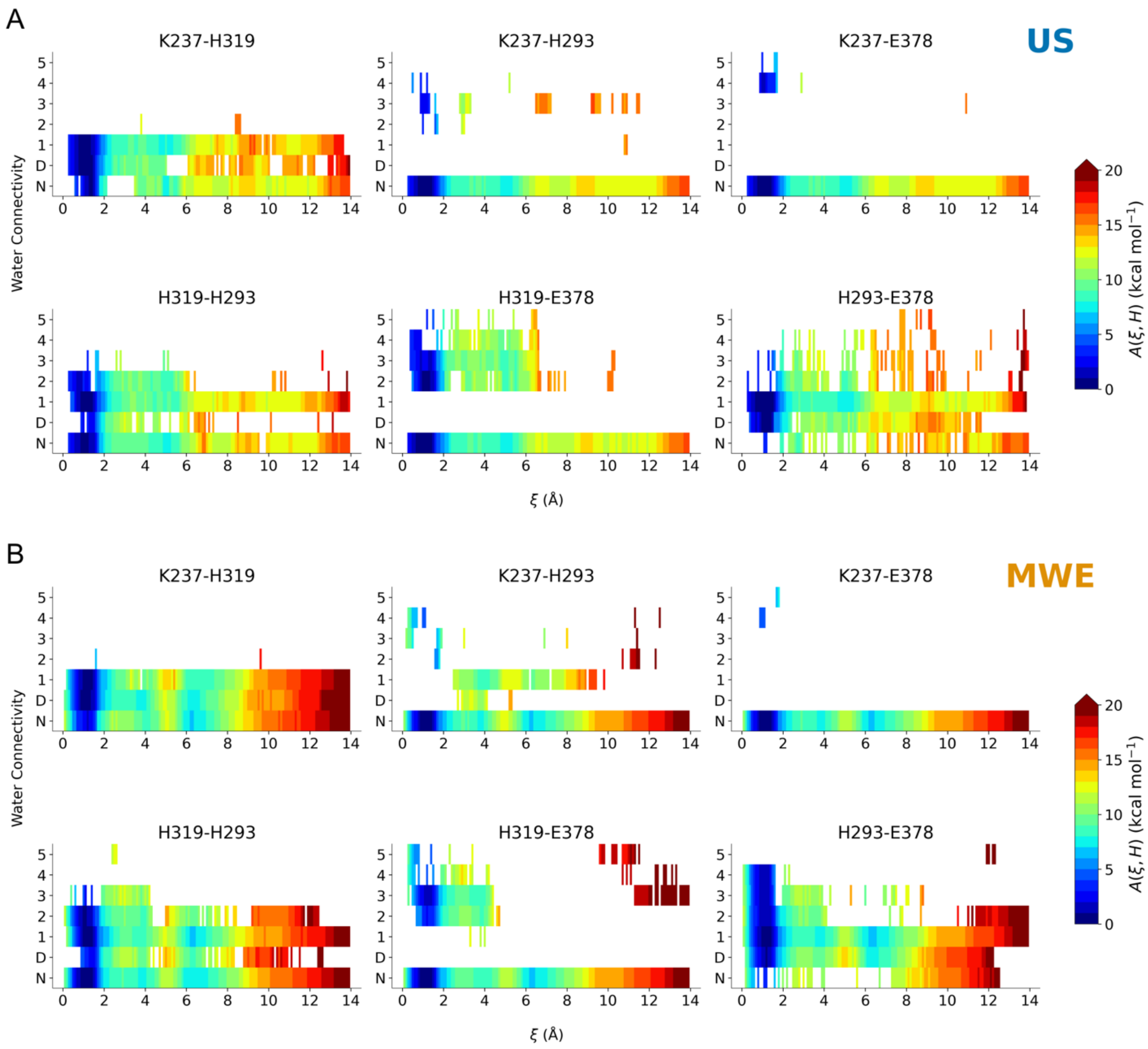


**Figure S12.** Sampled hydration states during proton transfer in the Complex I model as a function of mCEC-CV value. *N* stands for no water-mediated connection between two residues, *D* stands for a direct connection between two residues. (A) Water connectivity sampled with US. (B) Water connectivity sampled with MWE.

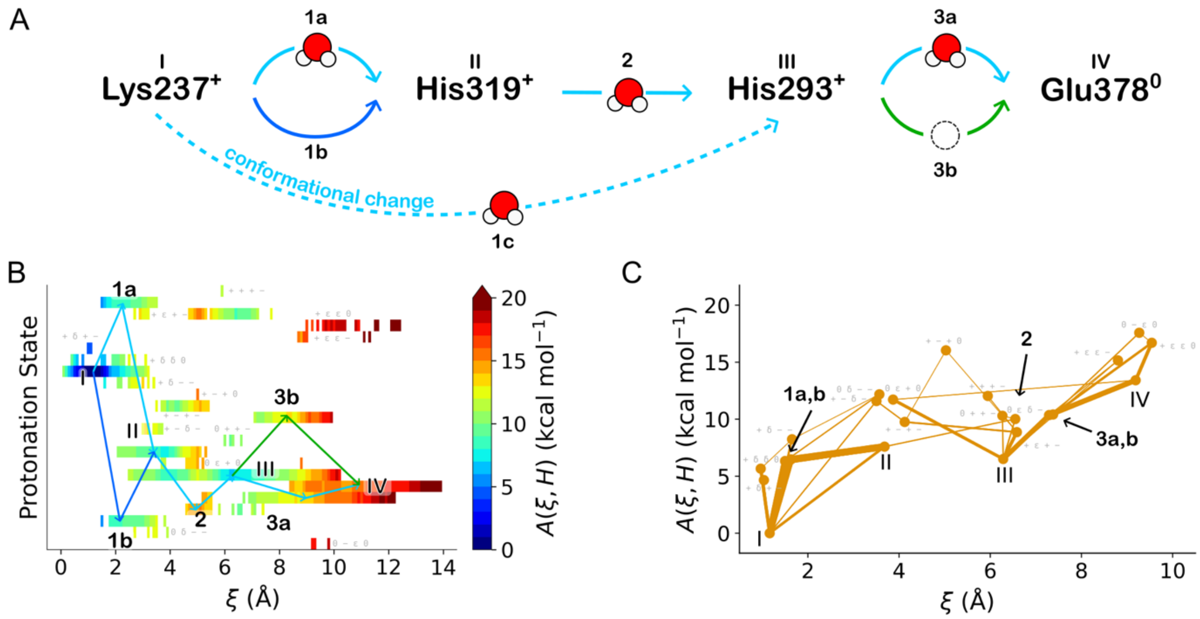


**Figure S13.** Sampling of protonation states during proton transfer in the Complex I model. (A) Reaction mechanism scheme sampled by MW-WTM-eABF. Water-mediated reaction steps are denoted by a cartoon water molecule, hole transfer is denoted by a dashed circle, direct protonation. Direct protonation is denoted by an unadorned arrow. (B) 2D-PMF along the mCEC-CV and the protonation states of the protein residues participating in the pT. Intermediates and arrows correspond to mechanism scheme in A. Additional protonation states are annotated in grey using the following scheme: protonation state of Lys237, His319, His293, and Glu378. (C) Protonation states plotted according to their weighted mCEC-CV average and their free energy. Transitions between two states during sampling are depicted as connections between the two respective states. Thicker lines denote more frequent transitions. Intermediate states are annotated using arrows.

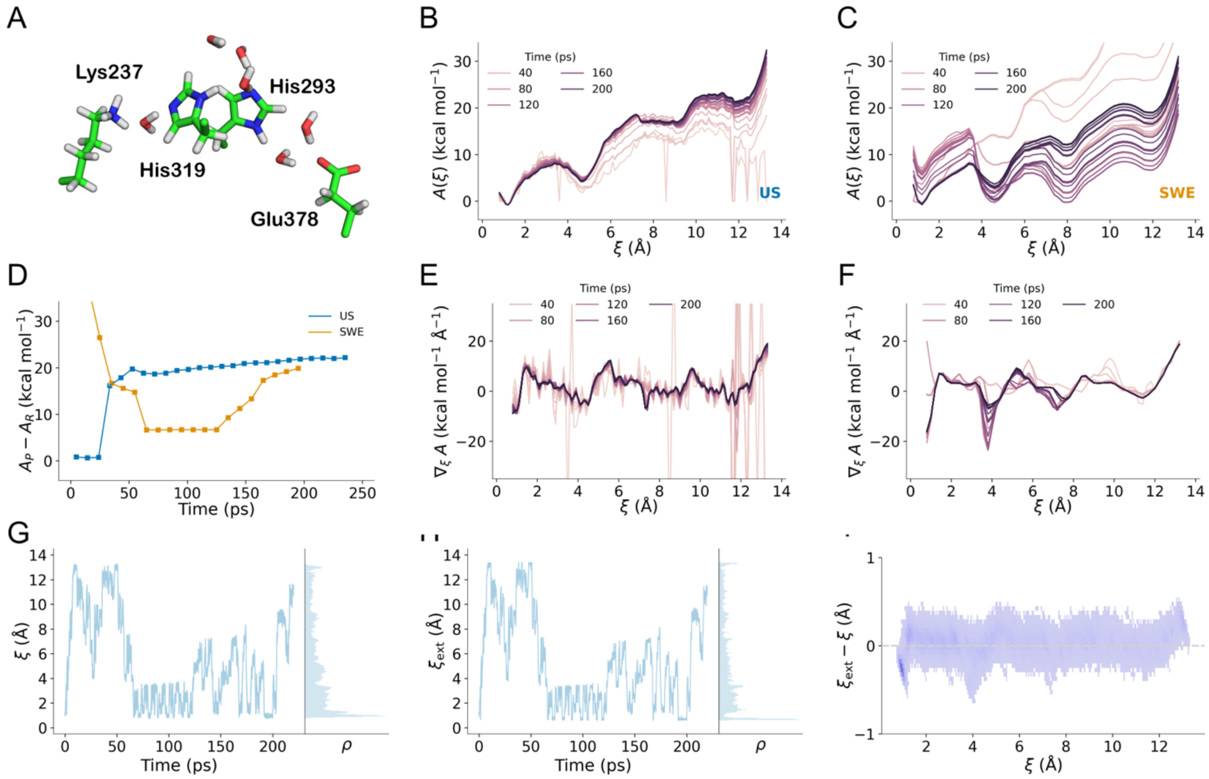


**Figure S14**. Sampling of proton transfer reactions in the Complex I modeled, using a smaller QM region and positional restraints. (A) Structure of the QM region. (B, C) Convergence of the PMF profile obtained with US and SW-WTM-eABF, respectively. (D) Convergence of PMF difference between reactant and product state. (E, F) Convergence of mean force profile obtained with US and SWE, respectively. (G) Sampling of the reaction coordinate using SWE. (H) Sampling of the extended variable reaction coordinate using SWE. (I) 2D histogram of the instantaneous difference between the extended system variable and the reaction coordinate as a function of the reaction coordinate.

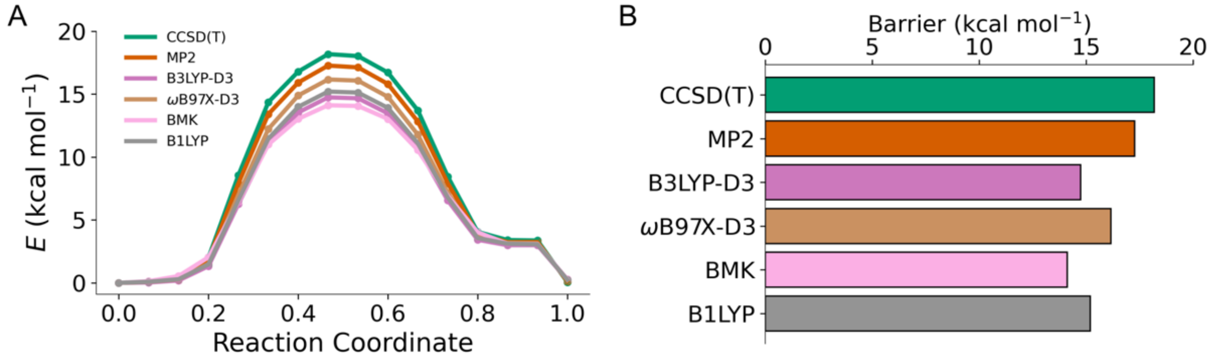


**Figure S15.** Benchmarking the proton transfer energetics for the His-water model (model 1). (A) Minimum energy profiles and (B) barriers at different theory levels. B3LYP-D3/def2-SVP geometries were used for estimation of electronic energies with the def2-TZVPPD basis set at the B3LYP-D3,[6-8] ωB97X-D3,[9] BMK,[10] B1LYP,[11] and compared against PNO-based CCSD(T) and MP2 level of theory.

**Table S1.** List of simulations.

| Simulation | Model | Reaction Coordinate | Sampling Algorithm | Sampling Time (ps) | | | Total Sampling Time (ps) |
|---|---|---|---|---|---|---|---|
| 1 | 1 | LC | US | 18 | × | 50 | 900 |
| 2 | | | MWE | 2 | × | 48 | 96 |
| 3 | | mCEC | US | 33 | × | 20 | 660 |
| 4 | | | MWE | 2 | × | 50 | 100 |
| 5 | | | MWE | 10 | × | 30 | 300 |
| 6 | 2 | mCEC | US | 60 | × | 5 | 300 |
| 7 | | | MWE | 12 | × | 25 | 300 |
| 8 | | | FUS | 12 | × | 10 | 120 |
| 9 | 3 | mCEC | US | 48 | × | 5 | 240 |
| 10 | | | SWE | 1 | × | 220 | 220 |

**Table S2.** Apparent barriers and activation free energies for Model 1 for different switching parameter $d_s$.

| $d_s$ (Å) | $m_{z^\ddagger}$ (a.m.u.) | $\Delta A^\ddagger$ (kcal mol$^{-1}$) | $A^\ddagger - A_R = \Delta A_{RP}^{\ddagger,app}$ (kcal mol$^{-1}$) |
|---|---|---|---|
| 0.03 | 0.017 | 8.4 | 10.5 |
| 0.05 | 0.026 | 8.5 | 10.4 |
| 0.10 | 0.049 | 8.7 | 10.0 |
| 0.20 | 0.086 | 8.6 | 9.6 |

### References


(1) Bakan, A.; Meireles, L. M.; Bahar, I. ProDy: Protein Dynamics Inferred from Theory and Experiments. *Bioinformatics* **2011**, *27* (11), 1575-1577. DOI: 10.1093/bioinformatics/btr168.
(2) Amadei, A.; Linssen, A. B. M.; Berendsen, H. J. C. Essential dynamics of proteins. *Proteins: Structure, Function, and Bioinformatics* **1993**, *17* (4), 412-425. DOI: 10.1002/prot.340170408.
(3) Pracht, P.; Bohle, F.; Grimme, S. Automated exploration of the low-energy chemical space with fast quantum chemical methods. *Physical Chemistry Chemical Physics* **2020**, *22* (14), 7169-7192. DOI: 10.1039/C9CP06869D.
(4) Bannwarth, C.; Ehlert, S.; Grimme, S. GFN2-xTB—An Accurate and Broadly Parametrized Self-Consistent Tight-Binding Quantum Chemical Method with Multipole Electrostatics and Density-Dependent Dispersion Contributions. *Journal of Chemical Theory and Computation* **2019**, *15* (3), 1652-1671. DOI: 10.1021/acs.jctc.8b01176.
(5) Dietschreit, J. C. B.; Diestler, D. J.; Hulm, A.; Ochsenfeld, C.; Gómez-Bombarelli, R. From free-energy profiles to activation free energies. *The Journal of Chemical Physics* **2022**, *157* (8). DOI: 10.1063/5.0102075.
(6) Becke, A. D. Density-functional exchange-energy approximation with correct asymptotic behavior. *Physical Review A* **1988**, *38* (6), 3098-3100. DOI: 10.1103/PhysRevA.38.3098.
(7) Lee, C.; Yang, W.; Parr, R. G. Development of the Colle-Salvetti correlation-energy formula into a functional of the electron density. *Phys. Rev. B* **1988**, *37* (2), 785-789. DOI: 10.1103/physrevb.37.785.
(8) Grimme, S.; Antony, J.; Ehrlich, S.; Krieg, H. A consistent and accurate ab initio parametrization of density functional dispersion correction (DFT-D) for the 94 elements H-Pu. *J. Chem. Phys.* **2010**, *132* (15), 154104. DOI: 10.1063/1.3382344.
(9) Chai, J.-D.; Head-Gordon, M. Long-range corrected hybrid density functionals with damped atom–atom dispersion corrections. *Physical Chemistry Chemical Physics* **2008**, *10* (44), 6615-6620. DOI: 10.1039/B810189B.
(10) Boese, A. D.; Martin, J. M. L. Development of density functionals for thermochemical kinetics. *The Journal of Chemical Physics* **2004**, *121* (8), 3405-3416. DOI: 10.1063/1.1774975.
(11) Adamo, C.; Barone, V. Toward reliable adiabatic connection models free from adjustable parameters. *Chemical Physics Letters* **1997**, *274* (1), 242-250. DOI: 10.1016/S0009-2614(97)00651-9.
